# Single-cell dissection of obesity-exercise axis in adipose-muscle tissues

**DOI:** 10.1101/2021.11.22.469622

**Authors:** Jiekun Yang, Maria Vamvini, Pasquale Nigro, Li-Lun Ho, Kiki Galani, Marcus Alvarez, Yosuke Tanigawa, Markku Laakso, Leandro Agudelo, Päivi Pajukanta, Roeland J. W. Middelbeek, Kevin Grove, Laurie J. Goodyear, Manolis Kellis

## Abstract

Regular physical exercise has long been recognized to reverse the effects of diet-induced obesity, but the molecular mechanisms mediating these multi-tissue beneficial effects remain uncharacterized. Here, we address this challenge by studying the opposing effects of exercise training and high-fat diet at single-cell, deconvolution and tissue-level resolutions across 3 metabolic tissues. We profile scRNA-seq in 204,883 cells, grouped into 53 distinct cell subtypes/states in 22 major cell types, from subcutaneous and visceral white adipose tissue (WAT), and skeletal muscle (SkM) in mice with diet and exercise training interventions. With a great number of mesenchymal stem cells (MSCs) profiled, we compared depot-specific adipose stem cell (ASC) states, and defined 7 distinct fibro-adipogenic progenitor (FAP) states in SkM including discovering and validating a novel CD140+/CD34+/SCA1-FAP population. Exercise- and obesity-regulated proportion, transcriptional and cell-cell interaction changes were most strongly pronounced in and centered around ASCs, FAPs, macrophages and T-cells. These changes reflected thermogenesis-vs-lipogenesis and hyperplasia-vs-hypertrophy shifts, clustered in pathways including extracellular matrix remodeling and circadian rhythm, and implicated complex single- and multi-tissue communication including training-associated shift of a cytokine from binding to its decoy receptor on ASCs to true receptor on M2 macrophages in vWAT. Overall, our work provides new insights on the metabolic protective effects of exercise training, uncovers a previously-underappreciated role of MSCs in mediating tissue-specific and multi-tissue effects, and serves as a model for multitissue single-cell analyses in physiologically complex and multifactorial traits exemplified by obesity and exercise training.

## Introduction

Obesity is a complex disease with genetic, environmental and behavioral origins that poses a public health problem of grave concern showing no signs of abating^1–3^. Key hallmarks of obesity include dysfunctional white adipose tissue (WAT) and chronic low-grade inflammation, which affect the function of multiple organs. Obese individuals are at significantly increased risk of type 2 diabetes (T2D), cardiovascular and cerebrovascular disease, and certain types of cancers, all leading causes of death in developed countries^4^. Regular exercise is known to improve metabolic function in numerous tissues, and to delay, prevent, or alleviate the effects of T2D, obesity, and cardiovascular disease^5^. Recent studies show that exercise-induced adaptations to WAT and skeletal muscle contribute to the benefits of exercise on health, with subcutaneous and visceral adipose depots exhibiting distinct adaptations to exercise training^6–10^.

WAT and skeletal muscle are highly heterogeneous and dynamic endocrine organs impacted by obesity and exercise. WAT comprises mature adipocytes (∼20-40% of the cellular content^11^) and the stromal vascular fraction (SVF), that latter consisting of multiple cell types including preadipocytes, mesenchymal stem cells, and a variety of immune cells^12^. Skeletal muscle consists of myofibers, surrounded by connective tissue, fibro-adipogenic progenitors (FAPs), and immune cells^13–16^. In WAT, obesity induces inflammation, alters cellular composition, and leads to a maladaptive WAT expansion to accommodate the excessive caloric intake^17^; by contrast, exercise training stimulates WAT beiging, alters key metabolic proteins, impacts tissue-tissue communication, and contributes to overall improved metabolism^6, 7, 18–21^. In skeletal muscle, obesity induces altered metabolism, intramuscular fat accumulation, increased insulin resistance, and impaired tissue regeneration and remodeling^22, 23^; by contrast, training induces improved regeneration potential, increased insulin sensitivity, and secretion of numerous autocrine, paracrine, or endocrine myokines mediating tissue crosstalk^24^. However, the genes and pathways mediating these opposing multi-tissue effects of exercise and obesity remain uncharacterized at single-cell resolution, despite their extraordinary clinical importance for the development of new treatments against T2D, metabolic diseases, and the obesity epidemic.

Here, we dissect the opposing effects of exercise training and diet-induced obesity in subcutaneous WAT (scWAT), visceral WAT (vWAT), and skeletal muscle at both tissue-level and single-cell resolution. We study tissue-level changes across 60 samples, deconvolved changes in the most abundant cell types, and single-cell changes in rarer cell types, by generating a single-cell atlas across 204,883 cells of 22 cell types and 53 cell states in obesity-exercise interventions across the three tissues (**Fig. 1a,b**). These include three distinct adipose stem cell (ASC) states in WAT depots showing shared and distinct pathways and upstream regulators, and six FAP states in skeletal muscle including a previously uncharacterized SCA1-subpopulation. Tissue-level analyses reveal exercise-induced upregulation of lipid metabolism and cellular respiration and downregulation of immune and extracellular matrix (ECM) pathways in all three tissues, and deconvolution reveals opposing changes in cell type abundance for exercise training vs. obesity. Single-cell analyses reveal changes in cellular states, cell type-specific gene expression/pathway/regulatory network changes, and cell-cell communication changes within and across tissues. In all three tissues, exercise training leads to down-regulation of ECM remodeling and up-regulation of circadian rhythmicity genes, driven primarily by mesenchymal stem cell (MSC) populations, and down-regulation of *Cdkn1a* and up-regulation of *Dbp* (two exercise-regulated genes) show consistent genetic and transcriptional effects in human tissues. Overall, our results provide a reference atlas of obesity-exercise single-cell changes in metabolic tissues, and reveal key roles for MSC in potentially mediating tissue-specific and inter-tissue communication changes in response to obesity and exercise interventions.

**Fig. 1:**
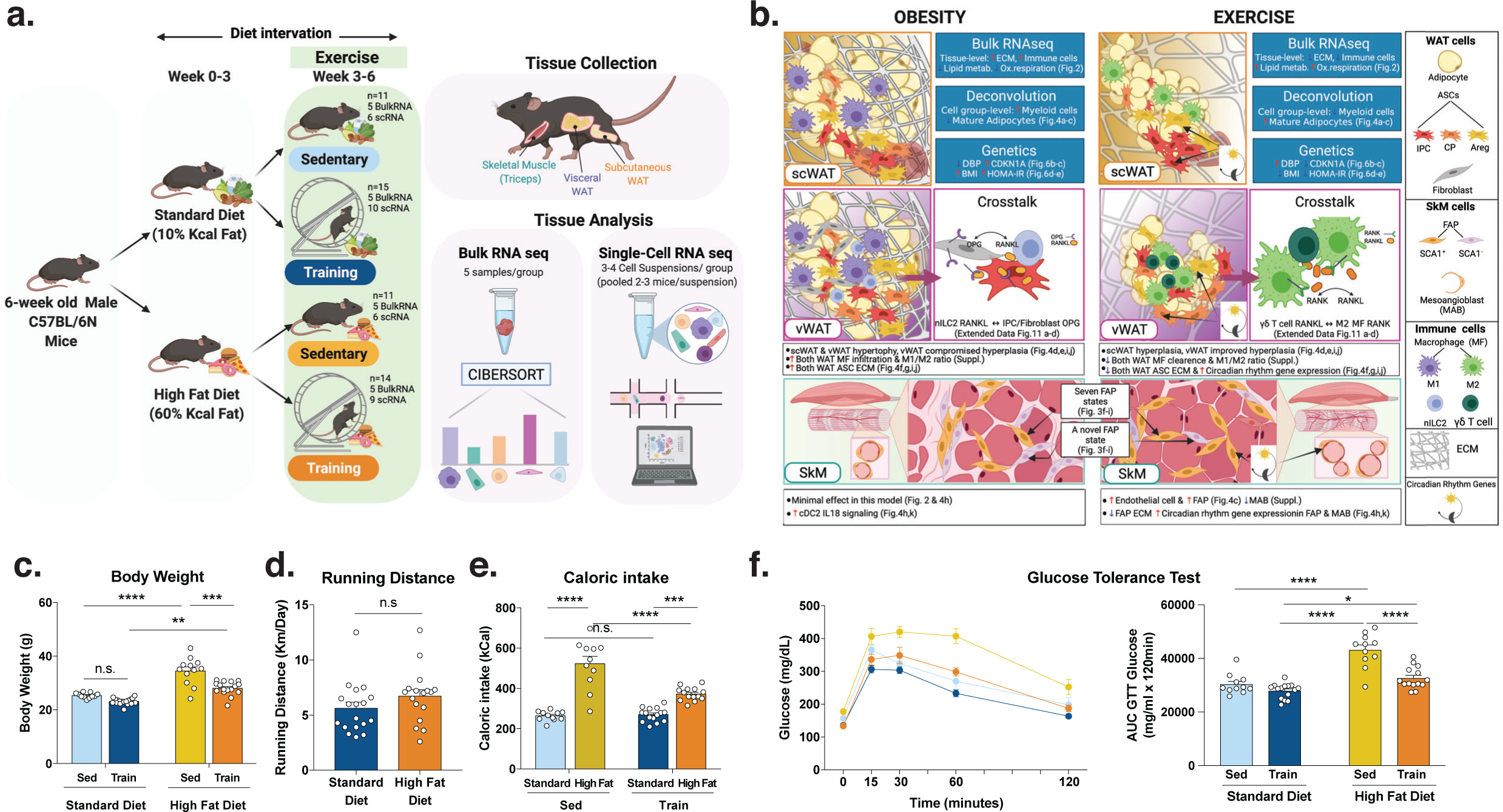
Study overview, highlighted results and phenotypic responses. **a,** We profiled bulk mRNA-seq and scRNA-seq in subcutaneous and visceral white adipose tissue (WAT), and skeletal muscle across 26 lean and 25 obese mice with 6-week diet and 3-week exercise interventions. **b,** Graphical representation and summary of highlighted results in this study. ECM, extracellular matrix. **c-f,** Body weight (**c**), running distance (**d**), caloric intake (**e**), and glucose tolerance test (GTT) result (**f**) in the four intervention groups. Statistical comparisons were performed using two way ANOVA followed by Tukey multiple comparison tests. P values are showns as *, < 0.05, ** < 0.01, *** < 0.001, **** < 0.0001. AUC, area under curve; n.s., not significant. Other abbreviations used in this figure appear in the Methods. Panels **a** and **b** were created with BioRender.com.

## Results

### Phenotypic response to diet-induced obesity and exercise, and profiling of three metabolic tissues

We subjected 6-week-old C57BL/6N male mice to diet-induced obesity (standard 10% vs. high-fat 60%, weeks 0-6) and exercise training (sedentary vs. voluntary wheel running, weeks 3-6) interventions (**Fig. 1a**, N=51 across 4 groups). Phenotypically, high-fat-diet induced, and exercise attenuated, weight gain and glucose intolerance (**Fig. 1c,f**); diet did not significantly impact running distance (**Fig. 1d**); and high-fat-diet animals consumed significantly more calories, especially when sedentary (**Fig. 1e**); each for both tissue-level and single-cell-level donors (**Supplementary Fig. 1a-e**).

We collected scWAT, vWAT and skeletal muscle (SkM) for both single-cell and tissue-level transcriptomic profiling after fasting (6-hour) and exercise wash-out (24-hour locked wheels) to investigate multi-tissue adaptations (**Fig. 1a**). For single-cell RNA-seq, we profiled 317,754 cells with 10X Genomics v3 across 42 libraries from 93 tissue samples (pooling 2-3 mice to obtain enough cells), capturing 6501 cells per library, 2025 genes per cell, and 45,421 reads per cell on average (**Supplementary Table 1**); after stringent quality control (QC), we report expression levels for a total of 17,341 genes across 204,883 cells (**Table 1, Extended Data Fig. 1a, Supplementary Table 1**). For tissue-level RNA-seq, we profiled 60 samples (4 groups x 5 mice x 3 tissues, no pooling), using 3′ Digital Gene Expression (DGE) RNA-seq. For both assays, the major drivers of variation were tissue, diet, and exercise, but not batch (**Extended Data Fig. 1b**).

**Table 1.** Number of cells and samples per tissue and phenotypic group used for analysis.

|  | Sedentary Std. Diet | Training Std. Diet | Sedentary High Fat Diet | Training High Fat Diet | Tissue Total |
| --- | --- | --- | --- | --- | --- |
| Subcutaneous White Adipose Tissue (scWAT) | 10,952 cells<br>3 samples* | 18,005 cells<br>4 samples | 12,783 cells<br>3 samples | 16,501 cells<br>3 samples | 58,241 cells<br>13 samples |
| Visceral White Adipose Tissue (vWAT) | 18,668 cells<br>3 samples | 20,869 cells<br>4 samples | 11,725 cells<br>3 samples | 27,598 cells<br>3 samples | 78,860 cells<br>13 samples |
| Skeletal Muscle (SkM) | 10,969 cells<br>3 samples | 21,969 cells<br>4 samples | 11,403 cells<br>3 samples | 23,441 cells<br>4 samples | 67,782 cells<br>14 samples |
| Total |  |  |  |  | 204,883 cells<br>40 samples |

\*each sample is a cell suspension

### Tissue-level gene and pathway alteration by obesity-exercise across three tissues

We found 1386 tissue-level differentially-expressed genes (DEGs) across all three tissues (568 in scWAT, 562 in vWAT, 256 in SkM; DESeq2-Negative-Binomial corrected P<0.05) and all three pairwise intervention comparisons for “obesity” (high-fat vs. standard diet, for sedentary), “training” (exercise training vs. sedentary, for standard diet), and “rescue” (exercise training vs. sedentary, for high-fat diet) (**Table 2**, **Supplementary Table 2**), and performed gene ontology (GO) pathway enrichment, and protein-protein interaction (PPI) analyses to reveal common and distinct biological processes across interventions and tissues (**Fig. 2a-d**).

**Fig. 2:**
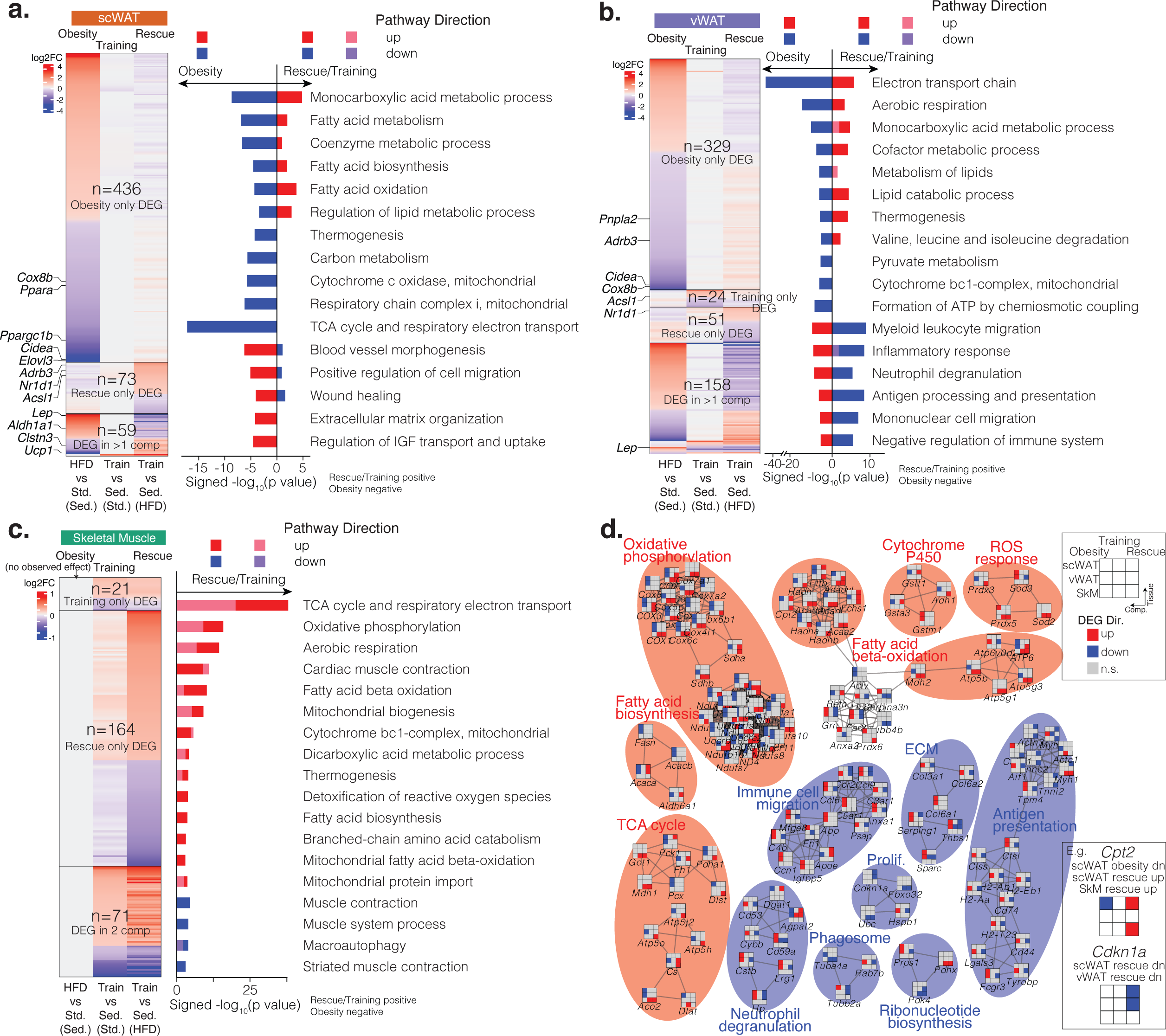
Tissue-level transcriptomic responses. **a-c,** Genes (heatmap) and pathways (bar plot) that are significantly differentially expressed and enriched across three comparisons: “obesity” (high-fat vs. standard diet under sedentary conditions), “training” (exercise training vs. sedentary under standard diet), and “rescue” (exercise training vs. sedentary under high-fat diet) in scWAT (**a**), vWAT (**b**), and skeletal muscle (**c**). The gene heat map is coloured by log_2_ fold change. The pathway bar plot is coloured by pathway direction in the three comparisons (red/pink: up-regulated, blue/purple, down-regulated). X-axis of the bar plot shows -log_10_p value with rescue/training pathways being positive, and obesity being negative. DEG, differentially expressed gene. **d,** Gene networks across selected DEGs from the three tissues that encode interacting proteins, clustered by protein-protein interactions with each cluster named by the most significantly enriched pathway. The 3-by-3 grid of each node (gene) is coloured by DEG direction in the three tissues (row) and comparisons (column). The cluster is coloured by DEG direction with exercise training. ECM, extracellular matrix; Prolif, proliferation; ROS, reactive oxygen species. Other abbreviations used in this figure appear in the Methods.

**Table 2.** Tissue-level differentially expressed genes (DEGs) with high-fat diet and exercise training interventions.

|  | Obesity (Sed. High-fat vs Sed. Std.) | Training (Train Std. vs Sed. Std.) | Rescue (Train High-fat vs Sed. High-fat) |  |
| --- | --- | --- | --- | --- |
| **Common in WAT** | **Up: 76;**<br>**Dn: 75** | **Up: 3;**<br>**Dn: 0** | **Up: 24;**<br>**Dn: 17** | **195 (176 unique genes)** |
| scWAT | Up: 269;<br>Dn: 223 | Up: 3;<br>Dn: 0 | Up: 70;<br>Dn: 62 | 627 (568 unique genes) |
| vWAT | Up: 237;<br>Dn: 241 | Up: 19;<br>Dn: 28 | Up: 72;<br>Dn: 133 | 730 (562 unique genes) |
| SkM | Up: 0;<br>Dn: 0 | Up: 64;<br>Dn: 28 | Up: 147;<br>Dn: 88 | 327 (256 unique genes) |
|  | 970 | 142 | 572 | 1684 (1386 unique genes) |

For the subset of DEGs identified in at least two comparisons in adipose tissues (59 in scWAT, 158 in vWAT), nearly all (94%-95%) stemmed from opposite changes in obesity vs. training/rescue (**Fig. 2a-b**). This anti-correlation held for all genes (not only DEGs), with obesity vs. rescue showing significant negative correlation in both depots (Pearson scWAT P<10^-10^, vWAT P<10^-16^) and obesity vs. training significantly anti-correlated in vWAT (Pearson P<10^-16^, **Extended Data Fig. 1c**). However, the subset of DEGs that met our stringent significance threshold in multiple interventions was small (10% and 28% in scWAT vWAT respectively), and most DEGs were significantly-different in only one comparison (77% and 59% in obesity only; 13% and 9% in rescue only; 0% and 4% in training only, in scWAT vWAT respectively).

Adipose-depot DEGs included both known and novel metabolism-associated genes. Known genes included obesity-upregulated and training/rescue-downregulated adiposity marker leptin^25^ and browning repressor *Aldh1a1*^26^, obesity-downregulated and training/rescue-upregulated browning/beiging driver *Ucp1*^26^, metabolism regulator *Pparg*, and multiple thermogenic and browning/beiging markers (*Cidea*, *Clstn3*, *Cox8b*, *Acsl1*, *Nr1d1*, *Adrb3*)^27, 28^. Novel genes included training/rescue-upregulated circadian rhythm regulators (*Dbp*, *Tef*, *Nr1d2*, *Per3*), and training/rescue down-regulated ECM remodelling genes (*Thbs1*, *Sparc*).

For SkM (**Fig. 2c**), rescue/training up-regulated 51 genes, including fast muscle fiber marker *Myh1*^29^, and down-regulated 20 genes, including fat metabolism repressor *Pdk4*^30^ and muscle-mass repressor *Asb2*^31^. While there was a limited transcriptional response to high fat diet intervention in SkM, an observation previously reported in short-term high fat diet interventions^32^, rescue showed significant anti-correlation with obesity across all genes (Pearson P<2.2x10^-16^, **Extended Data Fig. 1c**). Rescue also showed more DEGs than training (n=164 vs. 21), suggesting a stronger response of SkM to training under high-fat than standard diet. Similarly to WAT, training and rescue showed similar effect directions for most genes in SkM (**Extended Data Fig. 1c**).

These DEGs were enriched in tissue-specific pathways, including: (i) in scWAT, vessel morphogenesis and cell migration (obesity: up, rescue: down), and ECM organization and insulin-like growth factor (IGF) transport/uptake (obesity: up, **Fig. 2a**); (ii) in vWAT, immune pathways (obesity: up, rescue: down), which included myeloid leukocyte migration, neutrophil degranulation, and antigen processing and presentation (**Fig. 2b**); (iii) in SkM, muscle contraction processes and contraction-activated macro-autophagy (rescue: down) (**Fig. 2c**).

DEGs were also enriched in three tissue-shared pathway groups, including: (i) lipid metabolism, with both catabolic and anabolic fatty acid processes in all three tissues (obesity: down; rescue/training: up); (ii) respiration-related pathways (obesity: down, rescue/training: up), i.e., oxidative phosphorylation and detoxification of reactive oxygen species (ROS) in SkM, and cellular respiration and thermogenesis in WAT, consistent with obesity-associated hypoxia when mature adipocytes expand to oxygen-diffusion limit^33^; and (iii) basic metabolic processes (obesity: down, rescue: up), with monocarboxylic acid metabolic process in WAT, and branched-chain amino acid catabolism in SkM.

Across tissues and interventions, DEGs clustered into biologically-meaningful modules of interacting proteins (**Fig. 2d**). These include: (i) fatty acid biosynthesis/beta-oxidation/metabolism module (rescue/training up); (ii) cellular respiration modules (rescue/training up), including oxidative phosphorylation, TCA cycle, and ROS response; (iii) immune modules (rescue/training down), including antigen presentation, neutrophil degranulation, immune cell migration, phagosome-related genes; (iv) other modules (rescue/training down), including ECM-related genes, proliferation, and ribonucleotide biosynthesis. For example, Cpt2 involved in long-chain fatty acid oxidation in the mitochondria^34^ was down-regulated by obesity in scWAT and up-regulated by rescue in scWAT and SkM, and Cdkn1a involved in cellular senescence was down-regulated by rescue in scWAT and vWAT (**Fig. 2d**). Taken together, these tissue-level results reveal the specific genes and pathways that likely mediate the known beneficial effects of exercise training, specifically for improving fatty acid metabolism and cellular respiration across all three metabolic tissues, and repressing immune, ECM, proliferation and tissue-specific pathways.

### Single-cell atlas of metabolic tissues in obesity and exercise conditions

In addition to our tissue-level datasets, we also generated a single-cell atlas of 204,883 cells for obesity-exercise interactions across the three tissues and the four intervention groups (**Fig. 3a, Extended Data Fig. 2a-e**). To capture low-abundance cell types (e.g. ASCs in WAT, FAPs in SkM), we used a single-cell library preparation that enriched for stromal vascular fraction (SVF) instead of parenchymal cell types (e.g. mature adipocytes in WAT and muscle fibers in SkM) which are already well-captured by tissue-level studies, and included lymph nodes in scWAT to capture immune cells migrating between tissue and lymph nodes.

**Fig. 3:**
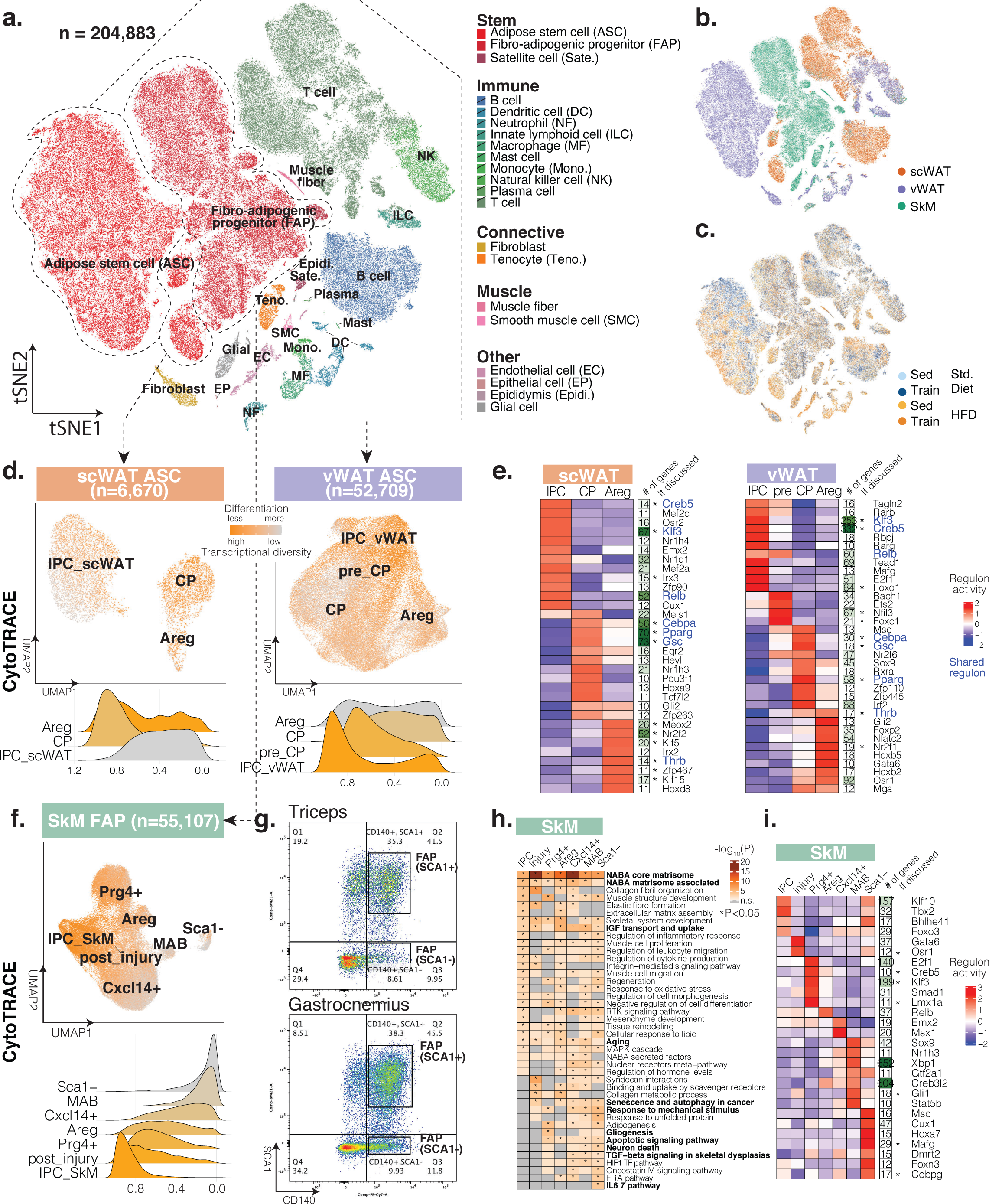
single-cell atlas and mesenchymal stem cell states characterization. **a,** Single-cell atlas of 204,883 cells across three tissues and four intervention groups. The tSNE plot is coloured by cell type (warm colours: non-immune cell types, cold colours: immune cell types). **b-c,** Single-cell atlas coloured by tissue (**b**) and intervention group (**c**). **d,** Re-clustering of ASCs in scWAT (left) and vWAT (right), coloured by CytoTRACE-predicted differentiation stage (orange: less differentiated, gray: more differentiated). Ridge plot of individual ASC states is colored similarly. **e.** Clustering of top ASC state-specific regulons (TF with the number of regulated genes as a separate heatmap column) in scWAT (left) and vWAT (right). Shared regulons across the two depots are coloured in blue, and regulons discussed in text are marked with asterisks. The heat map is scaled by column. **f,** Re-clustering of FAPs in SkM, coloured by CytoTRACE-predicted differentiation stage (orange: less differentiated, gray: more differentiated). Ridge plot of individual FAP states is colored similarly. **g.** FACS dot blot showing the sorting gates for Sca1+ and Sca1-FAPs from mouse triceps and gastrocnemius, with the percentages of the two populations labeled. **h-i.** Top pathways (**h**) and regulons (**i**) enriched in Sca1+ and Sca1-FAPs. The pathway heatmap is coloured by -log_10_p value. The regulon heatmap is coloured by activity score. A list of abbreviations used in this figure appear in the Methods.

We annotated 22 cell types using marker gene expression in cell clusters (hierarchical & density-based) of non-linear embeddings (tSNE, UMAP) for dimensionality-reduced data (top 50 PCs) (**Extended Data Fig. 1a & 2f-i, Supplementary Table 3, Methods**). These include: (i) 3 types of stem cells, including ASCs (MSCs in WAT), satellite cells (muscle stem cells in SkM), and FAPs (MSCs in SkM); (ii) 10 types of immune cells from both lymphoid and myeloid lineages; (iii) 2 types of connective cells, including tenocytes in SkM and fibroblasts primarily in vWAT; (iv) 2 types of muscle cells, including muscle fibers and smooth muscle cells; and (v) 4 additional cell types including endothelial, epithelial, epididymis, and glial cells (**Fig. 3a**). Within each cell type, our analysis revealed subclusters driven by tissue provenance (**Fig. 3b, Extended Data Fig. 3**, **Supplementary Table 4**) and to a lower degree by intervention group (**Fig. 3c**). Integration of our single-cell data with publicly available single-cell studies highlighted the strength of our analysis to capture lower abundance cell types as described earlier (**Supplementary Fig. 2**).

Subclustering revealed 42 cell subtypes/states for 11 (of the 22) cell types (**Extended Data Fig. 4**), distinguishing: (a) follicular vs. memory B-cells; (b) M1 vs. M2 macrophages; (c) T-cell subtypes including CD8 naive, CD8 cytotoxic, CD4 naive, CD4 memory, T regulatory (Treg), NKT, and naive (Cd27-) vs. memory (Cd27+) gamma-delta (Tgd); (d) dendritic cell (DC) subtypes conventional type 1, type 2, and monocyte-derived; (e) inflammatory vs. patrolling monocytes; (f) six tenocyte subtypes, including *Pdgfra*+ tendon stem/progenitor cells (TSPCs), pre_Dpp4+, *Dpp4+*, *Col22a1+*, *Pappa2+*, and *Scx-*; (g) vascular smooth muscle cells (SMCs) vs. SMC precursors vs. pericytes; (h) endothelial subtypes^35^ associated with large vessel vs. large artery vs. capillary vs. lymphatic vessel; (i) myelinating vs. non-myelinating glial (Schwann) cells; (j) three subtypes of ASCs in WAT, including interstitial progenitor cells (IPCs, marked by *Dpp4+*/*Pi16+*), committed pre-adipocytes (CPs, marked by *Icam1+*/*Fabp4+*), and adipogenesis regulatory cells (Aregs, defined^36, 37^ as CD142+/*Fmo2*+), and a fourth subtype specific to vWAT (pre_CP), lying between IPCs and CPs and expressing both *Dpp4* and *Icam1*; and (k) seven subtypes of FAPs, which we discuss below. Two-dimensional embeddings of each tissue showed clear distinctions of these cell subtypes (**Extended Data Fig. 5a-c**).

### Molecular signatures of depot-specific adipose stem cell (ASC) states

We evaluated WAT transcriptional diversity as a marker of differentiation potential^38^ (**Fig. 3d**). For vWAT, we confirmed that IPCs show increased transcriptional diversity, consistent with their earlier differentiation state^36^. In scWAT however, CPs and Aregs showed increased transcriptional diversity, which may be related to their beiging and de-differentiation potential^33^.

We sought co-expressed regulator/target-gene combinations (regulons) for each ASC subtype to gain insights on their gene-regulatory circuitries. Depot-shared regulons (**Fig. 3e, blue; Supplementary Table 6**) included: established regulons Klf3 and Creb5 for IPCs^39^; established regulons Cebpa, Pparg, Gsc for CPs^39^; and for Aregs, a regulon controlled by thyroid hormone receptor beta (Thrb), whose agonist (Resmetirom, MGL-3196) is in phase 3 clinical trial (NCT04951219) for Non-Alcoholic Fatty Liver Disease (NAFLD), suggesting Resmetirom might also act on Aregs in both WAT depots.

Depot- and state-specific regulons included: in scWAT, IPC-highest Irx3 (lowest in CPs), consistent with its early-adipocyte-differentiation role^40^; in vWAT, IPC-enriched Foxo1, consistent with its regulatory role in ASC differentiation^41^; in scWAT, Areg-enriched Nr2f2 and Meox2 regulons consistent with previous results^36, 37^, and scWAT-specific pro-adipogenic^36^ Klf5, Klf15, and Zfp467 regulons; in vWAT Aregs, human- and mouse-vWAT-specific^42^ (vs. scWAT) *Nr2f1*, suggesting Areg-enriched roles (instead of pan-vWAT); in vWAT pre-CPs, potential IPC-to-CP lineage restriction (commitment) regulators Foxc1, implicated in mesodermal commitment, and Nfil3, implicated in circadian rhythm and IGF-1 receptor signaling pathways.

### Seven distinct fibro-adipogenic progenitor (FAP) states and a novel FAP population in skeletal mus- cle

We classified the ∼55K FAPs into seven distinct cellular states using mouse and human markers^14–16^: (i) multipotent IPC_SkMs, sharing IPC markers with WAT; (ii) FAP_Cxcl14+, also found in other mouse studies^14^; (iii) FAP_Prg4+, also found in human^16^; (iv) adipogenesis-regulating FAP_Aregs similar to adipose tissue Aregs^15^; (v) FAP post injury representing an inflammatory post-injury state; (vi) Mesoangioblasts (MABs), marked by *Alpl*+; (vii) a previously-unreported, Sca1-subpopulation of FAPs, discussed in more detail below (**Extended Data Fig. 4k**). The first six subtypes were all positive for *Pdgfra*, *Cd34* and *Ly6a* (**Extended Data Fig. 5d**), all seven were detectable in previous single-cell data^13, 14^ (**Extended Data Fig. 5e**), and all lacked *Peg3* markers of PW1+/Pax7-interstitial cells (PIC)^43^ and *Abcg2* markers of muscle side population cells (SP)^44^ except for the population of post-injury FAPs, suggesting its overlap with PICs and SPs (**Extended Data Fig. 5f**).

Surprisingly, while FAPs are usually defined using interchangeable combinations of canonical markers *Pdgfra+*, *Cd34+,* and Sca1+, we found a new FAP population negative for Sca1 (**Extended Data Fig. 5d**)^45^. According to CytoTRACE differentiation prediction^38^, Sca1-FAPs showed a more differentiated state (less transcriptional diversity) than Sca1+ FAPs (**Fig. 3f**). We validated Sca1-FAPs for both triceps and gastrocnemius muscle by stringent fluorescence-activated cell sorting (FACS) and gating, and used qPCR to verify expression of Sca1+ marker genes (*Pdgfra*, *Ly6a, Dpt*) vs Sca1-markers (*Pdgfra*, Apoe), and lack of markers for endothelial cells (*Pecam1)*, tenocytes (*Scx)*, or glial cells (*Plp1)* (**Fig. 3g, Extended Data Fig. 5g,h, Supplementary Table 7**).

Marker genes for the seven FAP states were enriched in shared and unique pathways (**Fig. 3h, Supplementary Table 8**). Shared pathways across seven states included core matrisome, matrisome associated, IGF transport and uptake, and aging, suggesting FAP as a key contributor to ECM^46^, IGF signaling^47^, and aging^48^ in SkM. Unique pathways included: senescence, response to mechanical stimulus and apoptotic signaling pathways in all states except for IPC_SkM, suggesting specific FAP states mediating muscle regeneration in response to muscle damage^49^; gliogenesis and neuron death in FAP_Prg4+, MAB and Sca1-FAP, suggesting their involvement in neurodegeneration-mediated muscle atrophy^50^; IL6-mediated signaling pathway in Sca1-FAP, highlighting its potential to promote muscle atrophy and fibrosis^50^ and mediate muscle glucose uptake, increased insulin sensitivity, and increased fatty acid oxidation^24^.

FAP state-specific regulons supported their respective functions (**Fig. 3i**, **Supplementary Table 6**): in FAP_post_injury, Osr1 regulon, which marks adult FAPs activated by acute injury^51^; in FAP_Prg4+, Creb5, Klf3, and Lmx1a regulons, consistent with its enriched cellular response to lipid and adipogenesis pathways^52^ and suggesting its potential role in ectopic fat deposition in SkM; in MAB, Gli1 regulon, consistent with a recently-reported subpopulation of FAPs with higher clonogenicity and reduced adipogenic differentiation^53^; in Sca1-FAP, Mafg and Cebpg regulons, consistent with IL6-induced oxidative stress response^54^.

### Cell-type and cell-state proportion changes in obesity and exercise across three tissues

We also deconvolved^55^ our tissue-level data using independent single-cell maps^13, 56^ refined by manually curation (**Supplementary Fig. 3-5**)^56^. For scWAT and vWAT (**Supplementary Fig. 6a**), deconvolution captured 12 cell types: mature adipocytes (22-24% of cells, consistent with previous studies^11^), ASCs, smooth muscle cells, endothelial cells, pericytes, epithelial cells, glial cells, and five immune cell types (B-cell, NK cell, T-cell, macrophage, neutrophils). SkM samples (**Supplementary Fig. 6b**) deconvolved into 7 cell types: type II (fast) and type I (slow) myofibers (64-86% vs. 0-5% of nuclei, as expected for triceps^13^), mature adipocytes (5%), tenocytes, FAPs, endothelial cells, and macrophages. As expected, deconvolution captured primarily high-abundance cell types, but missed the vast majority of the 53 subtypes/states captured by single-cell profiling (**Supplementary Fig. 2**).

We used these tissue-deconvolution results to characterize cell type proportion changes in our three intervention comparisons: “obesity” (high-fat vs. standard diet, in sedentary), “training” (training vs. sedentary, in standard diet), and “rescue” (comparing training vs. sedentary, in high-fat diet). In both adipose tissues, obesity significantly decreased mature adipocytes and increased ASCs and myeloid cells (**Fig. 4a,b, Extended Data Fig. 6a,c**), consistent with increased adipocyte necrosis and macrophage infiltration in obesity-associated metabolic decline^33^; rescue reversed these effects in high-fat diet, even though training alone showed no effect (in standard diet). Histology staining in both WAT depots showed that obesity significantly enlarged mature adipocyte size (adipocyte hypertrophy) and increased tissue weight in obesity, which were both reversed by rescue (**Fig. 4d,e**). In SkM, training and rescue decreased type II (fast) myonuclei and increased FAPs, myeloid cells, and endothelial cells (**Fig. 4c**, **Extended Data Fig. 6g**), as in human^16^.

**Fig. 4:**
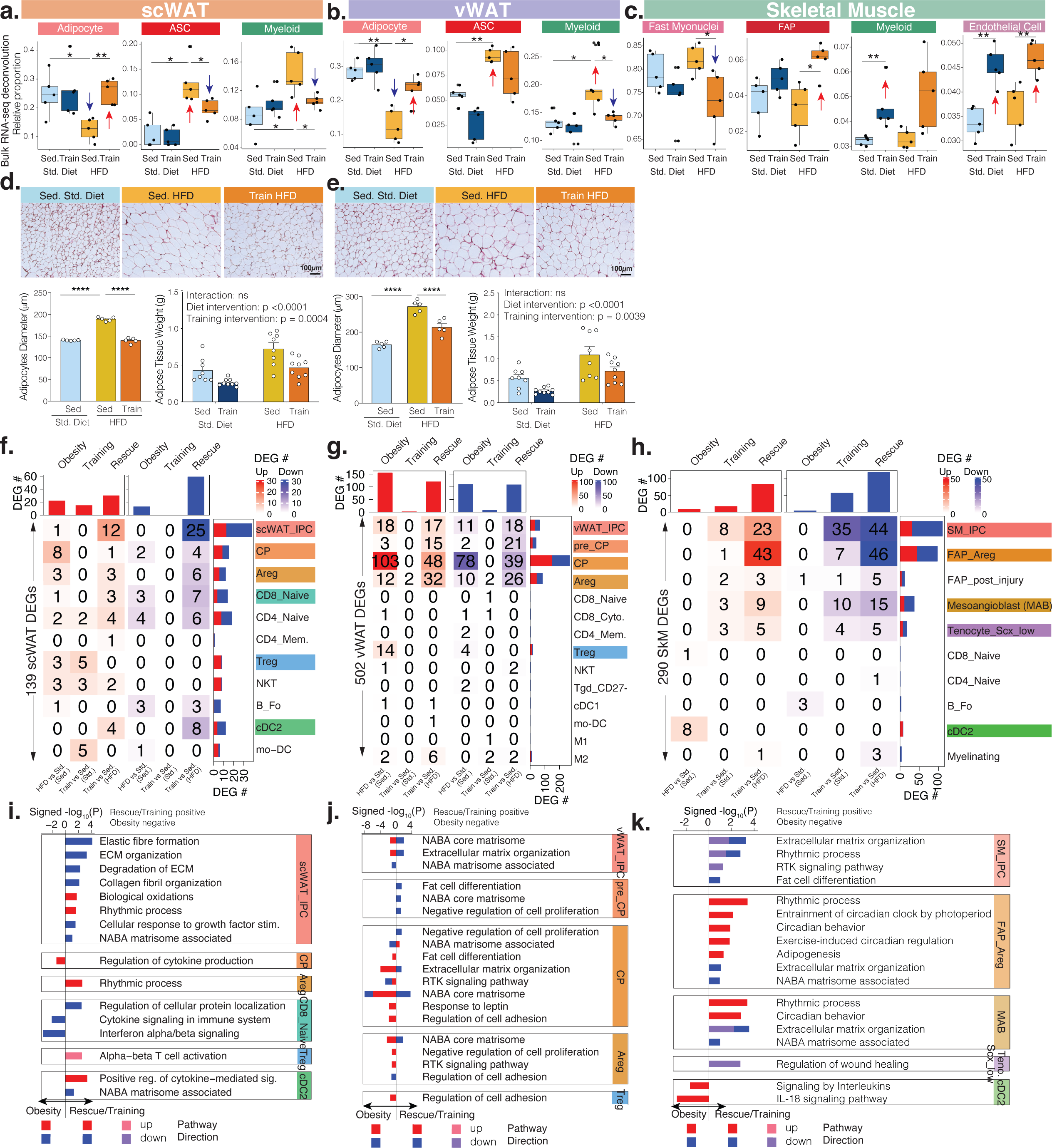
Single-cell-level proportion and transcriptomic responses across the three tissues. **a-c,** Sample-specific proportions of cell types across the four intervention groups in scWAT (**a**), vWAT (**b**), and SkM (**c**) after bulk RNA-seq data deconvolution. * p < 0.05, ** p < 0.01. **d-e,** Histology of scWAT (**d**) and vWAT (**e**) across three intervention groups, with bar plots below showing adipocyte diameter and adipose tissue weight changes across intervention groups. ****, p <0.0001. **f-h,** The number of cell state-specific DEGs (heatmap) that are up-regulated (red) or down-regulated (blue) in our three comparisons in scWAT (**f**), vWAT (**g**) and SkM (**h**). **i-k,** Pathways (bar plot) that are significantly enriched in cell state-specific DEGs across the three comparisons in scWAT (**i**), vWAT (**j**) and SkM (**k**). X-axis of the bar plot shows -log_10_p value with rescue/training pathways being positive, and obesity being negative. The bars are coloured by pathway direction in the three comparisons (red/pink: upregulated, blue/purple, down-regulated). DEG, differentially expressed gene. Other abbreviations used in this figure appear in the Methods.

We also used our single-cell results to annotate changes in cell subtype/state proportions (**Extended Data Fig. 6b,d**). In WAT across interventions, ASCs, macrophages and T-cells were the cell types with the most subtype/state proportion changes. In obesity, we found changes in vWAT specifically, with ASCs showing decreased CPs, which likely reflects both an increase in differentiation (into mature adipocytes, up-regulated fat cell differentiation in CPs with obesity, **Fig. 4j**) and decrease in CP proliferation. Rescue once more reversed the change, increasing CPs in vWAT (up-regulated cell proliferation in CPs with rescue, **Fig. 4j**). Integrating relative proportion with transcriptomic changes indicate obesity induced compromised hyperplasia with reduced CP proliferation, and rescue restored CP proliferation and improved adipogenesis in vWAT.

In addition to ASCs, immune cell subtypes also had relative proportion changes with obesity, training and rescue in vWAT (**Extended Data Fig. 6e,f**). Obesity both increased macrophage proportion and M1 (pro-inflammatory) vs. M2 (anti-inflammatory) ratio in the SVF of vWAT. Rescue restored the M1/M2 ratio toward baseline. In lymphoid lineage we found an increase of Tregs and a decrease of NKT cells in obesity, and these changes were reversed in rescue. Our findings are consistent with the current concept of the pathogenesis of obesity as it relates to the accumulation of Tregs in vWAT^57^, and a protective role of NKT cells against obesity via regulatory cytokines such as interleukin (IL)-4 and IL-10^58^. Furthermore, we observed similar changes of Tregs and NKT cells in training, suggesting that the beneficial effects of exercise training on lymphocytes occur regardless of diet. In SkM, within the MSC population, MABs significantly decreased in training and rescue, and showed a trend for an increase in obesity (**Extended Data Fig. 6h**). MABs primarily differentiate into myofibers, although they present a multipotent potential to differentiate into smooth muscle, adipocytes, or even osteocytes upon specific signaling^59^, suggesting high-fat diet and exercise training regulate lineage commitment of MABs.

### Exercise training regulates extracellular matrix (ECM) remodelling and circadian rhythm gene ex- pression across three tissues

We next used our single-cell data to infer cell-state-specific and cell-type-specific gene expression changes modulated by high-fat diet and exercise training interventions. In WAT (**Fig. 4f,g, Extended Data Fig. 7a, 8a, Supplementary Table 9**), we found 139 scWAT DEGs and 502 vWAT DEGs (3.6-fold more than scWAT) at the cell-state-level, affecting primarily ASCs (65+457 DEGs) followed by T-cells (49+29 DEGs) in both WAT depots. In ASCs, IPCs and CPs accounted for 57% and 59% of the ASC DEGs in scWAT and vWAT, respectively, indicating they are the ASC states most responsive to the two interventions. Between the two fat depots, obesity showed a 2.1-fold enrichment for vWAT DEGs (265 vs. 35), relative to expectation (2.5-fold more DEGs in vWAT). In vWAT, rescue reversed∼12% of ASC obesity DEGs, but 0% of T-cell DEGs, indicating that obesity-induced immune dysregulation is potentially longer-lasting compared to ASCs. In SkM, we identified 290 DEGs (13 obesity+74 training+203 rescue, **Fig. 4h, Extended Data Fig. 9a, Supplementary Table 9**). Rescue led to more single-cell DEGs than obesity and training, consistent with our tissue-level results. Sca1+ FAP showed the most DEGs, with IPC_SkM and FAP_Areg accounting for 81% of the Sca1+ FAP DEGs. Cell-type-level DEGs showed similar patterns (**Extended Data Fig. 7b,c, 8b,c, 9b,c**).

In scWAT, pathway analysis of the DEGs indicated that obesity up-regulated core matrisome and down-regulated type I interferon signaling and defense response, which, along with the vWAT increase in M1 macrophage population, suggests a shift from adaptive immune response to inflammation (**Fig. 4i, Extended Data Fig. 7, Supplementary Table 10**). By contrast, rescue enhanced ASC renewal and interferon signaling, and suppressed T-cell-dependent inflammation, likely mediated by increased Prostaglandin E2 levels, as evidenced by 1.5-fold decrease of prostaglandin degradation gene *Hpgd*. Consistently Prostaglandin E2 is regulated by cell-to-cell contact, helps maintain ASC self-renewal capacity in an autocrine manner, and enhances immunomodulatory potency^60^.

In vWAT, obesity up-regulated and rescue down-regulated two pathways that might mediate hypertrophy-to-hyperplasia adipocyte shifting induced by rescue (**Fig. 4j, Extended Data Fig. 8, Supplementary Table 10**): (a) TGFβ1 stimulus response, an anti-adipogenic inflammatory molecule secreted from hypertrophic, dysfunctional adipocytes and known to inhibit adipocyte differentiation in mice and humans^33^; and (b) ROS metabolic process, whose intracellular accumulation by mitochondrial respiration decreases preadipocyte differentiation^33^.

In SkM, obesity up-regulated and rescue down-regulated adipogenesis, immune signaling, and ECM pathways (**Fig. 4k, Extended Data Fig. 9, Supplementary Table 10**). Although FAPs mediate pro-myogenic signals that are critical for muscle homeostasis and regeneration, in myopathies and obesity, FAP adipogenesis has been reported to be released and causing fat infiltrates^61, 62^. Rescue altered adipogenesis pathways in two Sca1+ FAP states: in IPC_SkMs, rescue down-regulated adipocyte differentiation, consistent with increased Areg_FAPs (**Extended Data Fig. 6h**), which inhibit adipogenesis in SkM^15^; in Areg_FAPs however, rescue up-regulated adipogenesis pathways, including pro-adipogenic regulator *Klf15*, an Areg regulon in scWAT (but not in SkM). In the immune compartment, obesity up-regulated the IL-18 signaling pathway in conventional type 2 DCs, consistent with IL-18 increase in DCs in inflammatory myopathies^63^.

Across the three tissues, ECM-related pathways were up-regulated by obesity and down-regulated by training and rescue, specifically in scWAT IPCs, all the four states of vWAT ASCs (IPC, pre_CP, CP and Areg), and three Sca1+ FAP states (IPC_SkM, Areg_FAP and MAB, **Fig. 4i-k, Extended Data Fig. 7-9, Supplementary Table 10**). WAT ASCs have been explored to generate a dermal scaffold for wound healing, which was shown to produce even more ECM components (fibronectin, collagen, etc.) than fibroblasts^70^. ECM is a highly dynamic structure that is continuously modified in response to several stimuli and energy availability. Excessive deposition of ECM components has been observed with obesity in WAT to provide structural support to the enlarged mature adipocytes and at same time promote tissue fibrosis and hypoxia. In SkM, ECM is essential for muscle development, growth and repair and mechanical support during exercise. In addition, ECM provides a dynamic microenvironment by regulating cellular behavior and fate.

We observed another notable pathway commonly-regulated in all three tissues and only observed in single-cell data, circadian rhythm pathway. It was up-regulated by training and rescue, and enriched in MSCs, including FAPs in SkM and ASCs in WAT, consistent with adiposity up-regulation by circadian disruption in mouse preadipocytes^33^ (**Fig. 4i-k, Extended Data Fig. 7-9, Supplementary Table 10**). Upstream regulator analysis revealed *Dbp*, *Tef* and *Hlf*, three homologous PAR bZIP TFs sharing motif specificity^64^, as potential master regulators of these training-altered circadian pathways in specific MSC states (**Extended Data Fig. 10a,b**). The Dbp regulon was up-regulated by training/rescue in scWAT IPCs and vWAT CPs and Aregs. Tef was up-regulated by training/rescue and down-regulated by obesity in vWAT IPCs and Hlf was up-regulated by training/rescue and down-regulated by obesity in SkM Areg_FAPs.

DEGs identified using deconvolved tissue-level data were mostly regulated in opposite directions in obesity vs. training/rescue, and indicated similar pathways as detected in our bulk and single-cell data (**Extended Data Fig. 10c-e, Supplementary Table 11**). In scWAT, tissue-deconvolved mature adipocytes showed significant regulation of thermogenesis genes/beige markers: obesity down-regulated *Clstn3*, *Cox8b* and *Ppara*, and rescue up-regulated *Acsl1*, *Vegfa* and *Adrb3*. The enriched pathways in both fat depots include lipid metabolism and transport in mature adipocytes, ECM-related pathways and IGF transport and uptake in ASCs and myeloid cells, immune cell activation, regulation of inflammatory response and monocyte chemotaxis in lymphoid and myeloid cells. In SkM, 44 out of the 50 tissue-deconvolved DEGs were found within fast myonuclei, among which *Fbxo32* (highly expressed during muscle atrophy) and *Pdk4* (observed at the tissue level and discussed above) were down-regulated with both training and rescue. At the pathway level, training down-regulated tissue remodeling and up-regulated VEGFA-VEGFR2 signaling, consistent with VEGF rescues muscle loss in mice^65^, and rescue down-regulated actin filament-based process, muscle system process, regulation of chemotaxis, lipid modification, and cell growth.

### Exercise training reprograms within and cross-tissue cellular communication

Cells function collaboratively, communicating both directly and indirectly to drive physiological responses to interventions across tissues. Taking advantage of our high-resolution single-cell annotations, we used co-expression of interacting structure-based ligand-receptor pairs^66^ to predict pairwise cellular communication within and across tissues, and how they change in obesity, training, and rescue interventions at both cell-type and cell-state levels (**Fig. 5, Extended Data Fig. 11, Supplementary Fig. 7-10, Supplementary Table 12**).

**Fig. 5:**
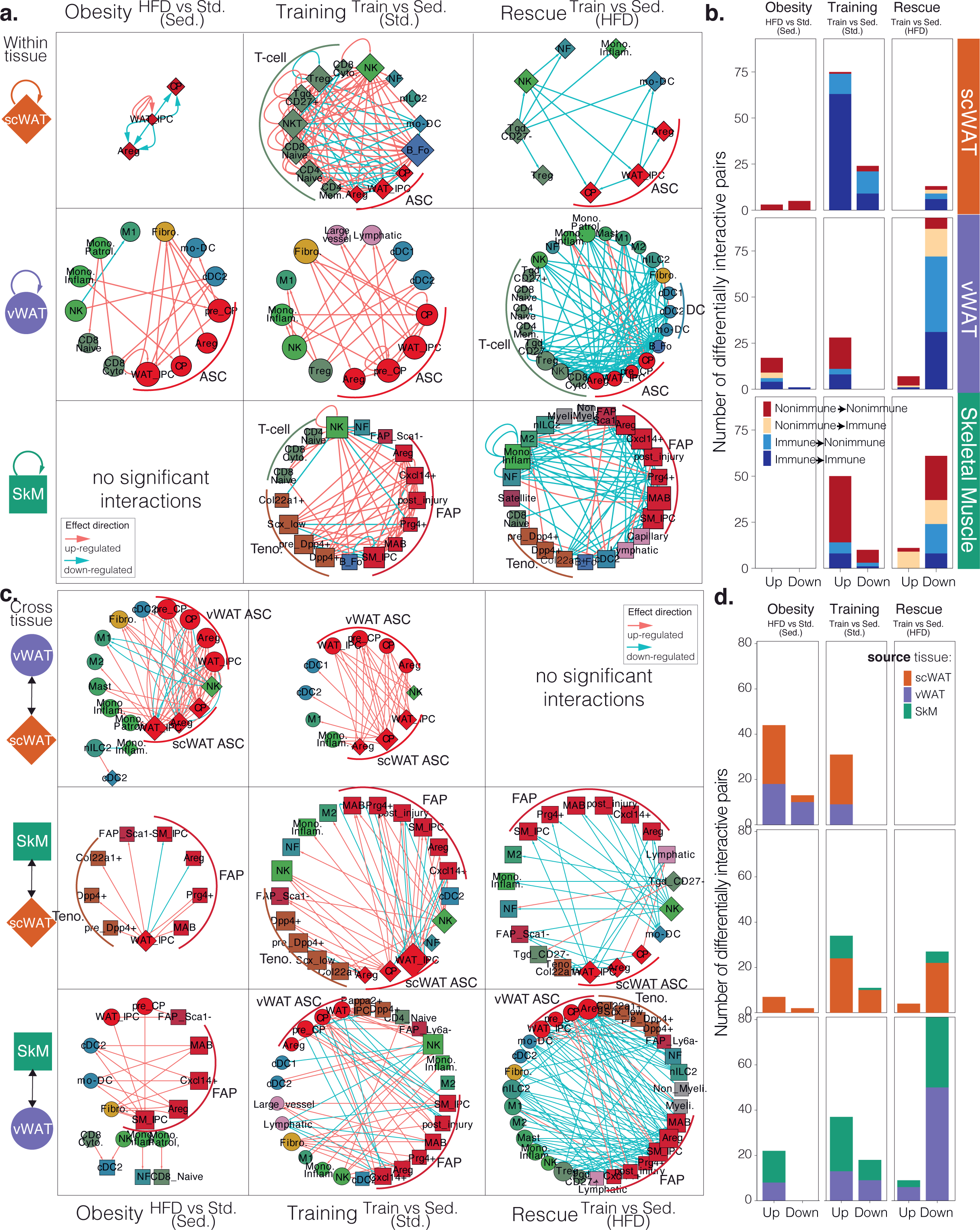
Within- and cross-tissue communication at cell-state level. **a,** Within-tissue ligand-receptor networks across the three tissues and three comparisons. Cell states (nodes) are shaped by tissue (diamond: scWAT, circle: vWAT, square: skeletal muscle) and sized by outdegree. Ligand-receptor interactions (edges) are directed, from ligand to receptor, and coloured by effect direction (pink: up-regulated, blue: down-regulated). **b,** The number of differentially interactive ligand-receptor pairs that are up- and down-regulated across the three tissues and three comparisons at cell-state level. Each bar is coloured by if the ligand or the receptor is from immune or non-immune cell state. **c,** Cross-tissue ligand-receptor networks across three pairs of tissues and three comparisons. The nodes and edges are formatted the same as in panel **a**. **d,** The number of differentially interactive ligand-receptor pairs that are up- and down-regulated across three pairs of tissues and three comparisons at cell-state level. Each bar is coloured by tissue source of the ligand. A list of abbreviations used in this figure appear in the Methods.

Within tissues, we observed MSCs (ASCs in WAT; FAPs in SkM) function as self-regulating and cross-regulating hubs of immune and non-immune cell types in individual intervention groups and in “obesity”, “training”, and “rescue” comparisons (**Fig. 5a,b, Extended Data Fig. 11a,b, Supplementary Fig. 7,8**).

In both fat depots, the sedentary high-fat diet group showed more interactions between ASCs and myeloid cells (DCs and macrophages). In vWAT specifically, fibroblast showed many interactions with ASCs and myeloid cells with exercise training or high-fat diet intervention. By comparing ranks of significant interactions overlapping intervention groups, we observed networks of differential interactions, including obesity up-regulated non-immune interactions (ASC autocrine regulation in scWAT, and ASC-fibroblast interactions in vWAT), training up-regulated interactions (immune-immune interactions in scWAT, ASC-endothelial-fibroblast interactions in vWAT, and FAP-tenocyte interactions in SkM), and rescue down-regulated immune to non-immune interactions (DC-T-ASC-fibroblast interactions in vWAT, and macrophage-FAP interactions in SkM). These observations implicate obesity mostly up-regulating non-immune interactions involving MSCs and rescue modulating immune-MSC interactions.

RANK (encoded by *Tnfrsf11a*, the receptor), RANKL (encoded by *Tnfsf11*, the ligand), and OPG (encoded by *Tnfrsf11b*, osteoprotegerin, the decoy receptor of RANKL) triad showed a particularly interesting expression change pattern in response to high-fat diet and exercise training interventions between fibroblasts and ILCs in vWAT. Obesity up-regulated and training and rescue down-regulated RANKL-OPG interaction (**Extended Fig. 11a**). Looking into expression patterns of the two genes across cell types/states in vWAT, we found OPG was mostly expressed in ASC IPCs and fibroblasts, RANKL was highly expressed in nILC2s and CD27-Tgds, and RANK was expressed mainly in M2 macrophages (**Extended Fig. 11c**). Obesity increased OPG expression in fibroblasts and IPCs and RANKL expression in nILC2 while training and rescue decreased OPG expression (**Extended Fig. 11d**). Interestingly, RANKL expression in CD27-Tgd and RANK expression in M2 macrophages showed an opposite trend of changes in obesity vs. training and rescue. This led us to hypothesize that high-fat diet promoted interaction between RANKL in nILC2 and the decoy receptor OPG in fibroblast and IPCs, and on the contrary, exercise training induced a shift from this relationship to interaction between RANKL from CD27-Tgd and RANK from M2 macrophage.

Across tissues, we identified biologically meaningful cellular interactions with differential activities in our three comparisons (**Fig. 5c,d, Extended Data Fig. 11e,f, Supplementary Fig. 9,10**). For example, training increased expression of *Tgfb2* (ligand) in scWAT ASCs and *Tgfbr3* (receptor) in SkM FAPs. We have shown in a previous study that TGF-β2, an exercise-induced adipokine from scWAT, is partially stimulated by lactate released from SkM during exercise, and its secretion improves glucose uptake in SkM^6^, for which our prediction suggested a mechanism via TGF-β2 interaction with Tgfbr3 in SkM, although the function of Tgfbr3 in FAP has not been studied yet^53^. Across all cross-tissue ligand-receptor pairs, obesity increased communication between fat depots, while training and rescue regulated SkM-WAT interactions as reported previously^24^. Although we restricted one of two interacting partners being secretable and the interaction being non-integrin for this analysis, without orthogonal data types such as metabolomics, lipidomics, and proteomics to provide additional evidence, the cross-tissue interactions we predicted should be interpreted with caution, especially for interactions between the two fat depots since they share most cell types.

### Genetics of two exercise-training candidate genes in two independent large-scale human studies

To evaluate the human relevance of our results, we tested anthropometric trait genetic associations in UK biobank^67^ and human metabolic tissue expression changes in Metabolic Syndrome in Men (METSIM) study participants^68^ for two commonly up/down-regulated genes across our three tissues by our tissue-level, tissue-deconvolution, and single-cell analyses (**Fig. 6a**).

**Fig. 6:**
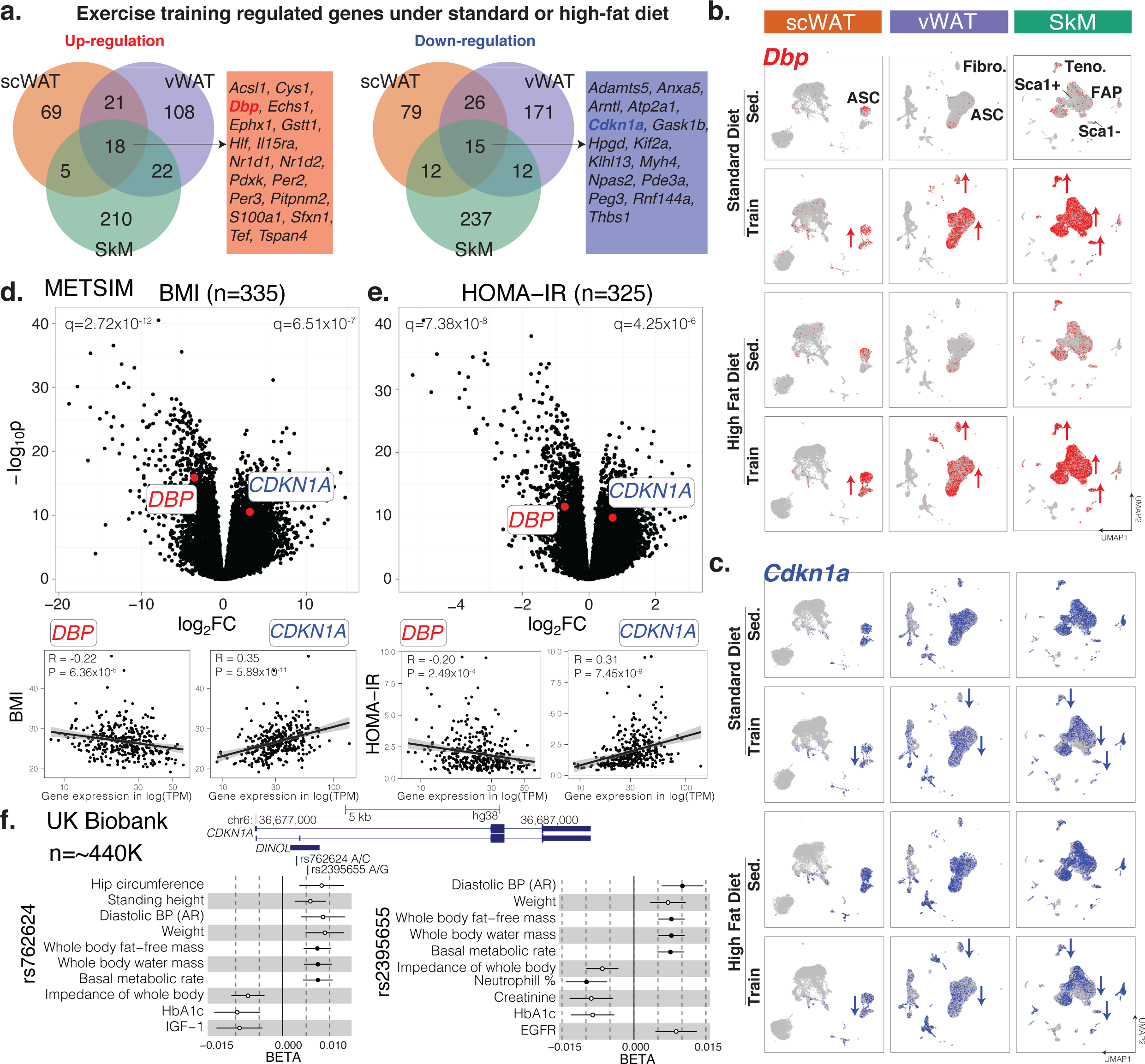
Two exercise-regulated genes (DBP and CDKN1A) in human. **a,** Overlap of up- and down-regulated genes by exercise training under standard or high-fat diet across the three tissues. Genes listed are regulated by exercise training in all three tissues. **b-c,** *Dbp* (**b**) and *CDKN1A* (**c**) expression across the three tissues and four intervention groups. Cell types with the most changes are labeled in the top panel. **d-e,** *DBP* and *CDKN1A* association with BMI (**d**) and HOMA-IR (**e**) in scWAT of METSIM subjects. Genes (dots in upper plots) and subjects (dots in lower plots) are plotted. HOMA-IR, homeostatic model assessment for insulin resistance. **f,** Association of two SNPs (rs762624 and rs2395655) in *CDKN1A* with anthropometric and metabolic traits in UK Biobank. The meta-analyzed PheWAS summary statistics (BETAs with standard errors, p < 1e-3) are shown. The filled circles are significant after correction (p < 1e-5). SNP, single nucleotide polymorphism; PheWAS, phenome-wide association study; BP: blood pressure. AR: automated reading. Other abbreviations used in this figure appear in the Methods.

Among 18 commonly-upregulated genes, we selected *Dbp*, a key regulator in diet/exercise-regulated circadian rhythm pathways, which is up-regulated in MSCs by exercise training across all three metabolic tissues (**Fig. 6b**), and whose homologs *Hlf* and *Tef* were also up-regulated. In human, *DBP* showed a significant negative transcriptional correlation with body mass index (BMI), homeostatic model assessment for insulin resistance (HOMA-IR), fasting insulin and glucose levels, C-reactive protein (CRP, a marker for inflammation), and waist-hip-ratio (WHR) adjusted for BMI, and a significant positive correlation with Matsuda insulin sensitivity index in METSIM (**Fig. 6d,e, Extended Data Fig. 12a-f**), confirming its obesity and exercise relevance in human phenotypes.

Among 15 commonly-downregulated genes, we selected cell proliferation and sentence regulator *Cdkn1a*, down-regulated by training/rescue primarily in MSCs in all three tissues (**Fig. 6c**), whose down-regulation was consistently found with diverse experimental (bulk and single-cell) and computational (different single-cell pipelines) methods. *CDKN1A* showed a significant positive correlation with BMI, HOMA-IR, fasting insulin levels, CRP, and WHR adjusted for BMI and a significant negative correlation with Matsuda insulin sensitivity index in METSIM (**Fig. 6d,e**, **Extended Data Fig. 12a-f**).

The CDKN1A genetic locus also contained two single nucleotide polymorphisms (SNPs), one intronic (rs762624) and one missense (rs2395655) in linkage disequilibrium (r^2^=0.49 in EUR), significantly associated with body weight, fat and fat-free mass, and energy expenditure (basal metabolic rate) in the UK Biobank (**Fig. 6f**). These variants were also significantly associated with hemoglobin A1C (HbA1c), a marker of long-term glucose control in diabetes. The minor allele of both SNPs (C for rs762624, G for rs2395655) showed protective associations with increased body fat-free mass, increased basal metabolic rate, and lower HbA1c. These SNPs are significant splicing quantitative trait loci (QTLs) for *CDKN1A* in the METSIM study, with the minor alleles increasing expression of a non-coding transcript isoform (long non-coding RNA, ENST00000462537.3, **Extended Data Fig. 12g,h**). These results suggest a potentially causal role for CDKN1A in human metabolic phenotypes, validating the human disease relevance of our results.

## Discussion

Our study, using complementary approaches of single cell, tissue-level, and tissue-deconvolution analyses, represents a large and comprehensive characterization of the molecular, gene-level, and pathway-level underpinnings of obesity-exercise changes and interactions in three key metabolic tissues (scWAT, vWAT and SkM). Our single-cell atlas with more than 200,000 cells and 53 annotated cell subtypes/states enabled identification and characterization of rare cell subtypes/states, specifically MSC populations in the three tissues, and revealed shared and distinct, known and novel biology of various cell types/subtypes/states within the three tissues in response to high-fat diet and exercise training (**Fig. 1b**).

Enrichment of rare cell types in our single-cell protocol led to large and high-quality MSC populations from all three tissues. In WAT, we uncovered IPCs, CPs, Aregs, and pre-CPs, a transitional state only observed in vWAT. We identified Foxc1 and Nfil3 as possible lineage-restricting upstream regulators transiently up-regulated in pre-CPs. Our gene regulator prediction for scWAT Aregs implicated regulators supporting previously reported discrepancy of this population of ASCs: some consistent with Schwalie *et al*.^37^ indicating an adipogenesis inhibitory function, and others supporting an adipogenic function reported by Merrick *et al*^36^. Interestingly, CPs and Aregs in scWAT showed higher transcriptional diversity than IPCs, which contrasts the differentiation hierarchy reported before^36^ and the hierarchy inferred in vWAT, but possibly provides transcriptional plasticity for beiging and dedifferentiation that are uniquely observed in scWAT^33^. We also revealed depot-specific signatures of the three ASC states including Irx3 and Foxo1 regulon enrichment in scWAT and vWAT IPCs, respectively.

In SkM, we defined seven distinct FAP states: IPC, Cxcl14 +, Prg4+, Areg, post-injury, MAB, and Sca1-. Areg_FAP was shown to inhibit adipogenesis^15^. Based on the enriched pathways and GRNs, Prg4+_FAP is likely to be another state involved in adipogenesis. Additionally, our identification and validation of Sca1-FAP, a novel MSC population in SkM, is reminiscent of a similar population reported in heart (cardiac fibrogenic SCA-1-cell)^69^, which were differentiated from PDGFRɑ+ SCA-1+ cells and showed a pathogenic role in post-myocardial infarction remodeling and arrhythmogenic cardiomyopathy^69^. Our Sca1-FAP was possibly differentiated from Sca1+ FAP, and it is a FAP state potentially responsive to exercise-induced IL6 production^24^.

Our single-cell data indicated that high-fat diet and exercise training modulate cell differentiation capability and proliferation rate, and gene expression of two pathways (ECM and circadian rhythm) within the three tissues, in a cell-state-specific manner. We identified diet-induced compromised **hyperplasia** in vWAT with decreased CP adipogenic capability and proliferation rate^36^, and exercise training-enhanced CP function which led to a shift from **hypertrophy**-to hyperplasia-dominant expansion in this depot. Orthogonal evidence (cell-state and deconvolved proportion and transcriptomics, and H&E staining) supports an exercise-driven shift from hypertrophy to hyperplasia in both adipose depots under high-fat diet.

At the pathway level, obesity up-regulated and exercise down-regulated **ECM**-related pathways in MSCs (major ECM contributors) across the three tissues. A recent report showed MSC-produced ECM components potentiate MSC response to differentiation stimuli through intracellular signaling pathway activation^71^. We observed the receptor tyrosine kinase (RTK) signaling pathway, a mediator of fundamental cellular and metabolic signaling pathways in response to growth factors, hormones, and cytokines^72^, tracks ECM changes in ASCs and FAPs. Interestingly, both the activity and substrate specificity of RTK signaling can be strongly influenced by ECM interactions, and controlled by ECM biochemistry and stiffness^73^. Besides MSCs, immune cells such as Tregs showed an up-regulation of cell adhesion with obesity potentially through ECM interactions. We highlight ECM as a dynamic entity remodeled by high-fat diet and exercise training to maintain tissue homeostasis by scaffolding and modulating different cell types.

One notable cell-state specific exercise-regulated pathway detected in our study was the **circadian** rhythm pathway, which was up-regulated by exercise training in particular MSC states of the three tissues. We identified Dbp and its homologs Tef and Hlf as potential master regulators of the up-regulated rhythmic pathway. Dbp is a PAR-domain TF with expression under circadian control, and emerging evidence suggests Dbp as an integral component of the peripheral circadian oscillator^74^. It is one of the circadian genes conserved among murine brown adipose tissue, scWAT, and vWAT^74^. Induction of *Dbp* was reported to ameliorate insulin sensitivity via direct binding to the promoter region of *Pparg*, driving mRNA expression of a splicing variant of *Pparg* (*Ppar-γ1sv*), and enhancing adipogenesis in preadipocytes from ob/ob mice vWAT^75^. Promoter activity of *Dbp* as reflected by histone H3 lysine 9 acetylation (H3K9ac) levels, and *Dbp* and *Pparg* expression in omental adipose tissue, were reported to be decreased in patients with T2D^76^. Our results suggest that exercise-induced circadian rhythm gene expression changes regulated by Dbp may contribute to adipogenesis of ASCs in WAT.

Single-cell data also empowered us to look into cell-cell communication within and across tissues. Our within-tissue analysis of vWAT highlighted an interesting protein triad, RANK-RANKL-OPG, with high-fat diet promoting interaction between RANKL in nILC2 and OPG in fibroblast and IPCs, and exercise training inducing a shift from this relationship to interaction between RANKL from CD27-Tgd and RANK from M2 macrophage. Although this triad has been mostly studied in bone and the immune system^77^, its role in WAT has been recently appreciated^78^: serum OPG is a biomarker for T2D, metabolic syndrome and obesity; global knockout of OPG in mice resulted in scWAT browning, resistance to high-fat diet-induced weight gain, and preserved glucose metabolism; and infusion of RANKL in wildtype mice led to scWAT beiging *in vivo*, and differentiation of scWAT-derived SVF and 3T3-L1 cells, but not mature white adipocytes, into beige adipose tissue^78^. Adipose tissue macrophages, the cell type expressing RANK receptor in our single-cell data, is a known player of beige adipogenesis as well^79^. Thus our results give prominence to an exercise-induced communication shift for RANKL secreted from naive Tgds to recruit M2 macrophages or re-polarize macrophages into an M2 state via RANK, for both their anti-inflammatory and beiging-eliciting effects.

Taking advantage of two independent large-scale human studies, we demonstrated human translational values of two exercise-training-regulated genes: higher expression of *DBP* and lower expression of *CDKN1A* in scWAT are associated with more desirable traits including lower BMI and HOMA-IR; minor alleles of two SNPs within *CDKN1A* (rs762624 and rs2395655) are associated with whole-body fat-free mass and HbA1c in ∼440,000 European subjects; these two variants are significant splicing-QTLs with the minor alleles increasing expression of a non-coding transcript isoform of *CDKN1A* in scWAT. Rs2395655 (missense) was associated with triglyceride measurement in ∼300,000 multi-ethnic participants of the Million Veteran Program^80^.

We envision several future directions for extending our study: (1) Surveying endocrine, cognitive, and other functions beyond the metabolic tissues surveyed here; (2) studying sex-specific adaptations^8^ beyond the male mice surveyed here; (3) adding two additional carolic intervention groups of high-caloric-intake exercise and low-carolic-intake no-exercise, to untangle the effects of caloric intake vs. exercise, as training impacted caloric intake in the high-fat-diet group; (4) distinguishing tissue-resident vs. circulating immune cells in scWAT by sampling lymph nodes vs. scWAT separately; (5) varying the length of high-fat diet and exercise, and the specific diet and exercise types; (6) testing mice of different ages; and (7) profiling epigenomic, splicing, proteomic, metabolomic, lipidomic, phosphorylation, and other molecular phenotypes.

Overall, our work provides a comprehensive and high-quality single-cell obesity-exercise interaction atlas in three metabolic tissues. We derived new insights on the metabolic protective effects of exercise training at single-cell transcriptome and interaction levels. We uncovered both a novel MSC population in SkM and a previously-underappreciated role of MSCs in potentially mediating tissue-specific and multi-tissue obesity and exercise training effects, with promise for new therapeutics development against obesity.

## Materials and methods

### Abbreviations

In many of the figure legends, abbreviations are used for tissues, intervention groups, cell types/subtypes/states. scWAT, subcutaneous white adipose tissue; vWAT, visceral white adipose tissue; SkM, skeletal muscle; sed, sedentary; train, exercise training; std. diet, standard diet; HFD, high-fat diet; ASC, adipose stem cell; FAP, fibro-adipogenic progenitor; sate, satellite cell; DC, dendritic cell; NF, neutrophil; ILC, innate lymphoid cell; MF, macrophage; mono, monocyte; NK, natural killer cell; teno, tenocyte; SMC, smooth muscle cell; EC, endothelial cell; EP, epithelial cell; epidi, epididymis; follicular, follicular B-cell; memory, memory B-cell; M1, M1 macrophage; M2, M2 macrophage; CD8 naive, CD8 naive T-cell; CD8 cyto, CD8 cytotoxic T-cell; CD4 naive, CD4 naive T-cell; CD4 memory, CD4 memory T-cell; Treg, regulatory T-cell; NKT, natural killer T-cell; Tgd CD27+, memory gamma-delta T-cell; Tgd CD27-, naive gamma-delta T-cell; cDC1, conventional type 1 dendritic cell; cDC2, conventional type 2 dendritic cell; moDC, monocyte-derived dendritic cell; inflammatory, inflammatory monocyte; patrolling, patrolling monocyte; TSPC, tendon stem/progenitor cells; pre_Dpp4+, transitional Dpp4+ tenocyte; Dpp4+, Dpp4+ tenocyte; Col22a1+, Col22a1+ tenocyte; Pappa2+, Pappa2+ tenocyte; Scx low, Scx-low tenocyte; vSMC, vascular smooth muscle cell; precursor, smooth muscle precursor cell; large vessel, large vessel endothelial cell; large artery, large artery endothelial cell; capillary, capillary endothelial cell; lymphatic, lymphatic vessel endothelial cell; myelinating, myelinating glial cell; non-myelinating, non-myelinating glial cell; IPC WAT, interstitial progenitor cell in white adipose tissue; pre_CP, transitional committed preadipocyte; CP, committed preadipocyte; Areg, adipogenesis regulatory/CD142+ cell; IPC SkM, interstitial progenitor cell in skeletal muscle; FAP Cxcl14+, Cxcl14+ fibro-adipogenic progenitor; FAP Prg4+, Prg4+ fibro-adipogenic progenitor; FAP Areg, adipogenesis regulatory/CD142+ fibro-adipogenic progenitor; FAP post injury, post-injury/inflammatory fibro-adipogenic progenitor; MAB, mesoangioblast; FAP_Sca1-, Sca1-fibro-adipogenic progenitor.

### Mouse cohort

All experiments were conducted following NIH guidelines and protocols were approved by the Institutional Animal Care and Use Committee at Joslin Diabetes Center in Boston, MA. C57BL/6N mice were purchased from the Charles River Laboratories and were housed in singular cages at room temperature (23°c) on a 12 h/12 h light/dark cycle in an AAALAC-approved animal facility at Joslin Diabetes Center. We used 6-week-old male mice for this experiment. For the first 3 weeks, all mice were sedentary, and half of the mice were fed a chow standard diet (10% kcal fat; 9F5020-LabDiet, Pharma-Serv,Inc.) and the other half a high-fat diet (60% kcal fat; (9F5020-LabDiet, PharmaServ,Inc.) ad libitum. High-fat feeding was used as a robust model for the development of impaired glucose tolerance. At the start of week 4, mice were further divided into sedentary and training groups, resulting in four groups: sedentary chow-fed; exercise training chow-fed; sedentary high-fat fed; and exercise training high-fat fed. The mouse cohort consisted of 60 mice: 12 in each sedentary group (12 chow;12 high-fat) and 18 in each training group (18 chow;18 high-fat). Exercise training was done by housing mice in individual cages containing a running wheel. Mice had free access to the wheel at all times, and running distance was recorded daily. Sedentary mice were individually housed in standard cages. All mice had body weights measured every two days. After 21 days, mice underwent a glucose tolerance test (GTT) after a 12-hour fast. Seven days later (day 28), the wheels of the trained mice were locked. Twenty-four hours later, following a 6-hour fast, mice were anesthetized with 5% v/v Isoflurane (NDC 60307-120, Piramal Healthcare) using EZ-150C anesthesia machine (E-Z Systems, Inc.) and blood was drawn by heart puncture. Perigonadal visceral (vWAT) and inguinal subcutaneous white adipose tissue (scWAT) and triceps muscle were rapidly dissected and were either snap frozen or processed fresh to generate cell suspension.

### Bulk mRNA sequencing

For the whole-tissue RNAseq, also known as bulkRNAseq, we euthanized 5 mice per group and harvested inguinal and perigonadal WAT as well as triceps. These tissues were snap frozen in liquid nitrogen immediately after collection. Total RNA was extracted at the Goodyear lab using an RNA extraction kit (Direct-zol™ RNA MiniPrep, Zymo Research). 10ng of total RNA quantified and quality assessed by Advanced Analytical Fragment Analyzer was used for library preparation on Tecan Evo150. 3′ DGE-custom primers 3V6NEXT-bmc#1-12 were added to a final concentration of 1 uM. (5’-/5Bi-osg/ACACTCTTTCCCTACACGACGCTCTTCCGATCT[BC6]N10T30VN-3’ where 5Biosg = 5’ biotin, [BC6] = 6bp barcode specific to each sample/well, N10 = Unique Molecular Identifiers, Integrated DNA technologies), to generate two subpools of 15 samples each. After addition of the oligonucleotides, Maxima H Minus RT was added per manufacturer’s recommendations with Template-Switching oligo 5V6NEXT (10uM, [5V6NEXT : 5’-iCiGiCACACTCTTTCCCTACACGACGCrGrGrG-3’ where iC: iso-dC, iG: iso-dG, rG: RNA G]) followed by incubation at 42℃ for 90’ and inactivation at 80℃ for 10’. Following the template switching reaction, cDNA from 12 wells containing unique well identifiers were pooled together and cleaned using RNA Ampure beads at 1.0X. cDNA was eluted with 17 ul of water followed by digestion with Exonuclease I at 37℃ for 30 minutes, and inactivated at 80℃ for 20 minutes. Second strand synthesis and PCR amplification was done by adding the Advantage 2 Polymerase Mix (Clontech) and the SINGV6 primer (10 pmol, Integrated DNA Technologies 5’-/5Biosg/ACAC-TCTTTCCCTACACGACGC-3’) directly to the exonuclease reaction. Eight cycles of PCR were performed followed by clean up using regular SPRI beads at 0.6X, and eluted with 20ul of elution buffer (Qiagen). Successful amplification of cDNA was confirmed using the Fragment Analyzer.

Illumina libraries were then produced using standard Nextera tagmentation substituting P5NEXTPT5-bmc primer (25μM, Integrated DNA Technologies, (5’-AATGATACGGCGACCACCGAGATCTACAC TCTTTCCCTACACGACGCTCTTCCG*A*T*C*T*-3’ where * = phosphorothioate bonds.) in place of the normal N500 primer. Final libraries were cleaned using SPRI beads at 0.7X and quantified using the Fragment Analyzer and qPCR before being loaded for sequencing using the Hiseq 2000 (Illumina, Inc) in 50bp single-end mode at the BioMicro Center at MIT.

### Single-cell RNA sequencing

Tissues from 2-3 mice were pooled to achieve >1x10^6^ analyzable cells (final sample size per group= 3-4). Fresh tissues were enzymatically digested and dissociated according to tissue dissociation kit protocols (adipose tissue Cat No 130-105-808, skeletal muscle Cat No 130-098-305 by Miltenyi) using the gentleMACS™ Dissociator (Miltenyi). Dissociated tissues were filtered, centrifuged, and the isolated cells were resuspended in 0.1% BSA in PBS and immediately processed for the generation of single-cell RNA (scRNA) libraries using the droplet-based RNA sequencing technology. Briefly, 5000-6000 cells were profiled per sample using the Chromium Single Cell 3’ RNA reagent kit v3 according to the 10X Genomics protocol. The generated cDNA libraries were indexed, pooled, and sequenced in three batches using the NovaSeq 6000 S2 system and reagent kits (100 cycles) (Illumina, Inc) at the BioMicro Center Core at MIT.

### Analysis of bulk mRNA-seq data

#### Pre-processing and DE analysis

Six FASTQ files for each sample were concatenated for read deduplication using unique molecular identifiers (UMIs). We then ran Salmon 0.14.2^81^ to quantify the number of unique reads for each transcript against Ensembl version 98 mouse transcripts. The transcript level information was summarized to the gene-level using R package tximport^82^. We then clustered all the samples across three tissues unbiasedly to observe potential batch effects and sample outliers.

Next for each tissue, genes with a count greater than 10 in all the samples were retained, and differential gene expression analysis for our three comparisons was carried out using R package DESeq2^83^.

We used Independent Hypothesis Weighting (IHW)^84^ to adjust p values and adaptive shrinkage estimator (ashr)^85^ to adjust fold changes from DESeq2 results. We called significant DEGs at an adjusted p value cutoff of 0.05 (**Supplementary Table 2**).

#### Deconvolution

We performed deconvolution on bulk mRNA-seq data using CIBERSORTx^86^ and in reference to two publicly available datasets^13, 56^: one is a scRNA-seq data in scWAT and the other is a snRNA-seq data in skeletal muscle. We integrated and re-annotated the scWAT scRNA-seq dataset as shown in **Supplementary Fig. 3 and 4**, and confirmed original cell type annotation for the skeletal muscle snRNA-seq dataset (**Supplementary Fig. 5**). We ran CIBERSORTx using default parameters for all three analysis modules, including creating signature matrices using the two reference datasets, imputing cell fractions, and sample-level gene expression using our bulk data. We calculated expression changes for genes with imputed expression levels in the three comparisons using the Wilcoxon rank sum test.

### scRNA-seq data analysis

#### Pre-processing, clustering and annotation

Gene count matrices for each single-cell sample were generated by aligning reads to the mm10 genome (refdata-gex-mm10-2020-A) using 10X Genomics Cell Ranger software v4.0.0 (**Supplementary Table 1**). We clustered pseudo-bulk profiles of individual single-cell samples to determine potential batch effects, and excluded one low-quality sample based on unbiased clustering results. Then for each sample, we removed ambient RNA contamination using SoupX^87^ with a fixed contamination fraction of 20%. The 20% fixed threshold performed the best compared to no ambient RNA removal, automatic removal implemented in SoupX, 10% and 15% fixed thresholds, and threshold estimated using hemoglobin genes, to reach a desirable de-contaminated visualization and keep the most number of cells. We then excluded low-quality cells using four QC metrics: (i) number of genes with non-zero expression fewer than 500; (ii) number of UMIs fewer than 200 or more than 6000; (iii) percentage of reads mapping to mitochondrial genes more than 10; and (iv) number of reads mapped to Mki67 more than 0. We removed potential cell doublets using Doublet-Finder^88^ with default parameters and 3.1% homotypic doublet proportion estimation based on statistics published by 10X Genomics. Next we integrated all the 41 samples across three tissues together for an atlas, and samples belonging to a single tissue together for tissue-specific maps. Integration was done without any batch correction using Seurat v3^89^. We used “sctransform” in Seurat for data normalization, performed principal component analysis (PCA) to obtain the first 50 PCs, used the 50 PCs to build community, and clustered the cells using both a graph-based clustering approach as implemented in Seurat v3 and a density-based clustering approach in R package dbscan^90^. Visualization of the tissue- and atlas-level datasets was through non-linear dimensional reduction techniques such as tSNE and UMAP. We adjusted processing steps for tSNE in reference to this paper^91^. We then annotated cell clusters using SciBet^92^, SingleR^93^, and cell type-specific markers from *Tabula Muris*^94^ and tissue-focused studies in the field^14, 36, 95, 96^. For unknown cell type subclusters like Sca1-FAP, we identified cell type-specific markers using the “FindMarkers’’ function in Seurat with an expression cutoff of 25% in either of the two tested populations. We further subsetted each cell type and performed sub-clustering within the cell type to identify cell subtypes/states. We annotated identified cell subtypes/states referencing markers from the literature (**Supplementary Table 3**).

#### DE analysis

We performed cell-type- and cell-state-specific differential expression (DE) analysis on “pseudo-bulk” profiles, generated by summing counts together for all cells with the same combination of cell type/state and sample. This approach leverages the resolution offered by single-cell technologies to define cell types/states, and combines it with the statistical rigor of existing methods for DE analysis involving a small number of samples^97^. The DE analysis was performed using quasi-likelihood (QL) methods from the edgeR package^98^. We removed cell type/state and sample combinations containing fewer than 10 cells. Cell-type/state-specific DEGs were determined using an FDR cutoff of 0.05 (**Supplementary Table 9**).

#### Cellular communicatio

For within- and cross-tissue communication prediction, we exported gene-by-cell count matrices and cell type/state assignment for each cell as two input files for CellPhoneDB “statistical analysis”^66^. CellPhoneDB is a publicly available repository of curated receptors, ligands and their interactions with the advantage of taking subunit architecture into consideration. We then imported CellPhoneDB results into R, merged interactions identified in each sample, and compared rank and mean values for all the interacting partners of an interaction in our three defined comparisons (“obe-sity”, “training”, and “rescue”). Specifically for cross-tissue interactions, we forced ligands to be secretable and interactions to be non-integrin. We then derived log2 fold changes using mean value, and calculated statistical significance on ranks using the Wilcoxon rank sum test implemented in base R. Interactions with a nominal p value of 0.1 were plotted using RCy3 package^99^ and Cytoscape (**Supplementary Table 12**).

#### Gene regulatory network and CytoTRACE

We inferred per-sample GRNs using SCENIC with GRN-Boost implementation in Python^100^, and detected regulons with differential activities between intervention groups using the Wilcoxon rank sum test. We estimated differentiation stages of ASCs and FAPs using the script provided with the CytoTRACE framework^38^. All the QC steps and analysis performed on single-cell data were illustrated and summarized in **Extended Data Fig. 1a**.

### Histology

Representative samples of scWAT and vWAT were fixed with 3.7% formaldehyde for 24hrs and then stored in 70% ethanol at 4°C. Five-micrometer-thick tissue sections were stained with hematoxylin and eosin (Richard Allan Scientific) and relative adipocyte size was estimated at 20 fold magnification of 5 random fields using an inverted fluorescence microscope (IX51Olympus). CellProfiler 3.0 (http://cellpro-filer.org) was used for the automatic measurement.

### FACS-based cell isolation, RNA isolation and quantitative PCR

Single cell suspension passed through 40 um (FisherBrand) and dead cell removal MS column (Miltenyi Biotec) were stained with anti-CD45 BV650 antibody (1:100, Biolegend), anti-CD34 PE antibody (1:100, Biolegend), anti-CD140 PE-Cy7 (1;100, Biolegend), anti-SCA1 BV421 (1:100, Biolegend) in PBS containing 0.1% BSA at 4°C for 30 min. After 3 times of washing, Cytox green (1:100, Invitrogen) was added as a dead cell marker. Live CD140+ SCA1+ and CD140+SCA-cells from CD45-CD34+ populations were sorted on Aria II (BD biosciences), and the RNA were isolated using Trizol in combination with miRNeasy kit (Qiagen). 15 ul of elution buffer was used to elute RNA, and we performed qRT-PCR using RNA to Ct kit on Quant Studio 7 (Thermo Scientific) to detect gene differentially expressed in CD140+SCA1+ and CD140+SCA1-.

### METSIM and UK Biobank analysis

#### METSIM RNA-seq

All participants provided informed consent and the study was approved by the ethics committee of the University of Eastern Finland. The METSIM cohort consists of 10,197 Finnish men with detailed metabolic phenotyping^68^. Among these, we analyzed 335 participants with RNA-seq data from subcutaneous adipose tissue biopsies. Reads were mapped using STAR v2.5.2b^101^ to the GRCh38 genome with Gencode^102^ v26 as a transcriptome annotation. Gene read counts were calculated using FeatureCounts. We performed transcriptome-wide differential expression for insulin, glucose, Homa-IR, C-reactive protein (CRP), free fatty acids (FFA), Matsuda index, BMI, and waist-hip ratio adjusted for BMI (WHRadjBMI). All phenotypes except WHRadjBMI were log-transformed to induce an approximate normal distribution. To improve power, we included RIN, the first PC, and sequencing batch as covariates. Normalization factors for library size were calculated using Trimmed Mean of M-values (TMM). To perform DE, we used edgeR v3.22.5^98^ with the quasi-likelihoods to fit the models and obtain p-values. P-values were adjusted for multiple testing using FDR.

#### METSIM isoform QTL

To estimate isoform transcript abundance, we ran Kallisto^103^ using Gencode v26 as a transcriptome reference. Isoform QTLs were detected with FastQTL^104^. Isoform transcripts per million (TPM) estimates from Kallisto were rank transformed to a standard normal distribution. FastQTL was run using RIN, batch, and the first 35 PCs as covariates. To determine the number of PCs, a QTL analysis was run on chromosome 21 with successively larger numbers of PCs as covariates. We selected 35 as this roughly maximized the number of isoform QTLs. We extracted nominal p-values and corrected for multiple testing as follows. For each isoform, we corrected SNP-isoform p-values using Benjamini-Hochberg. Then, we corrected the q-values for the number of isoforms tested using Bonferroni.

#### UK Biobank

To assess the phenome-wide associations of the genetic variants in *Dbp* and *Cdkn1a* across anthropometric and metabolic traits, we investigated GWAS summary statistics across 8 variants in the two candidate genes and 108 traits in UK Biobank. Briefly, we focused on meta-analyzed GWAS summary statistics on directly genotyped arrays^105^ across a total of 451,354 individuals consisting of white British (n = 337,129), non-British white (n = 44,632), African (n = 6,497), South Asian (n = 7,831), East Asian (n = 1,704), semi-related (n = 44,632), and admixed (n = 28,656) individuals, defined from a combination of genotype PCs and self-reported ancestry as described elsewhere^106, 107^.

The association summary statistics was visualized with R ‘ggforestplot’ package (https://nightin-galehealth.github.io/ggforestplot/index.html) and is available as a part of Global Biobank Engine^108^.

### Other computational analyses and data processing remarks

Enrichment analysis was performed using the web server Metascape with default parameters^109^. All the heatmaps were generated using R package ComplexHeatmap^110^. All computational analyses were performed using R version 3.4.0. All Wilcoxon rank sum tests were unpaired and two-sided. All two way ANOVA followed by Tukey multiple comparison tests were generated in GraphPad Prism v9. All box plots were generated and displayed in R, using the geom_boxplot() function with default parameters.

The median value is indicated with a black line, and a coloured box (hinges) is drawn between the 1st and 3rd quartiles (interquartile range, IQR). The whiskers correspond to no further than 1.5 x IQR from the hinge and outliers are omitted. All bar plots for phenotype analysis were generated and displayed in Prism, which display mean values as centres and the standard deviation as error bars. All included microscopy images are representative.

## Data availability

All raw and processed bulk mRNA-seq and scRNA-seq data have been uploaded in the GEO database (https://www.ncbi.nlm.nih.gov/gds) with the accession numbers xxx and xxx. We also provide an interactive data and analysis browser for all the data at xxx.

## Code availability

Analysis code is available at xxx.

## Acknowledgements

We thank Minna Kaikkonen-Määttä for advice and feedback on the work. We thank Stuart Levine for generating high-quality data and technical support. We thank Carlos Castorena and Angie Bookout for discussions and feedback on the high-fat diet and exercise training-regulated genes. We thank Karen Mohlke for analysis advice on the METSIM cohort. We thank Charlie Whittaker for help with bulk 3′ DGE data processing. We thank Ashley Renfro, Nicholas Carbone and Julio Mantero for experimental support. We thank all the members of the Kellis lab at MIT, the Goodyear lab at Joslin Diabetes Center, and the Kaikkonen-Määttä lab at University of Eastern Finland for feedback on the work. This work was supported by Novo Nordisk Research Center, Seattle, WA, USA. J.Y. was supported by the National Institutes of Health (NIH) National Institute of Diabetes and Digestive and Kidney Diseases (NIDDK) grant T32-DK110919. M.V. was supported by the NIH NIDDK grants T32-DK007260 and F32-DK126432. R.J.W.M. was supported by the NIH NIDDK grant K23-DK114550. L.J.G. was supported by R01DK099511, and 5P30-DK-36836. M.K. was supported by U24DK112331, U24HG009446, and UG3NS115064.

## Author Contributions

This study was designed and directed by L.J.G. and M.K. M.V. and P. N. performed the mouse protocol. M.V., P.N., L.H., and K.Ga. collected the tissues. K.Ga. performed the scRNA-seq experiment. J.Y. performed data processing and computational analysis. M.A. performed analysis in the METSIM study. M.L., and P.P. designed and directed the METSIM study. Y.T. performed analysis in UK Biobank. M.V., P.N., and L.H. performed validation experiments. M.K., L.A., R.J.W.M., and K.Gr. provided scientific feedback. J.Y., M.V., P.N., L.J.G., and M.K. wrote the manuscript.

---

**Extended Data Fig. 1:**
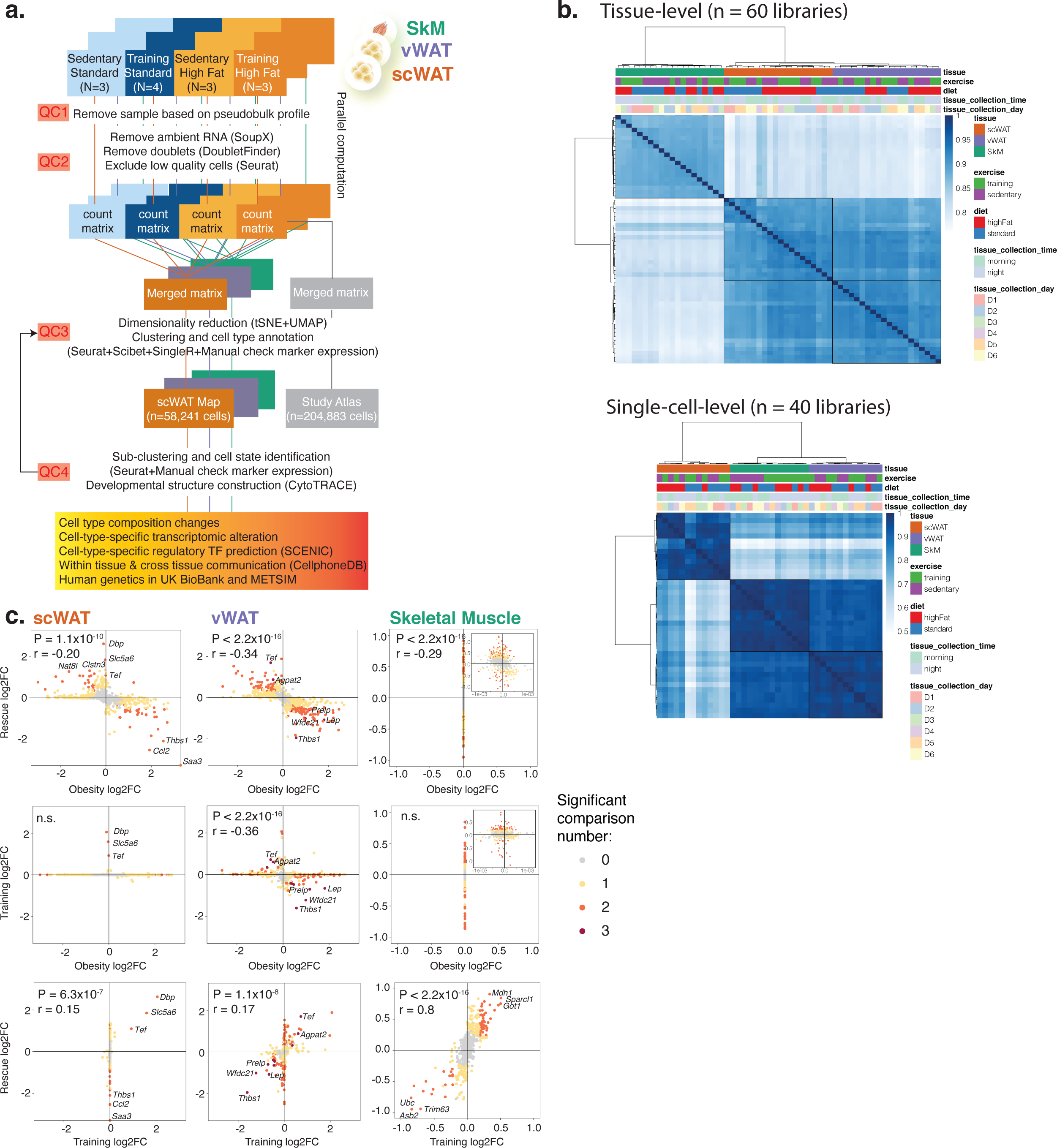
Single-cell analysis pipeline, sample clustering, and cross-comparison tissue-level transcriptomic change correlation. **a,** Graphical representation of single-cell analysis pipeline and the four QC steps taken. **b,** Unbiased clustering of tissue- and single-cell-level samples with variables that may introduce batch effects colored on the top. The variables include tissue type, exercise group, diet group, and tissue collection time and day. **c,** Global transcriptomic change correlation across “obesity”, “training”, and “rescue” comparisons in scWAT (left), vWAT (middle), and skeletal muscle. The P and r values were obtained from correlation tests. FC, fold change. Other abbreviations used in this figure appear in the Methods.

**Extended Data Fig. 2:**
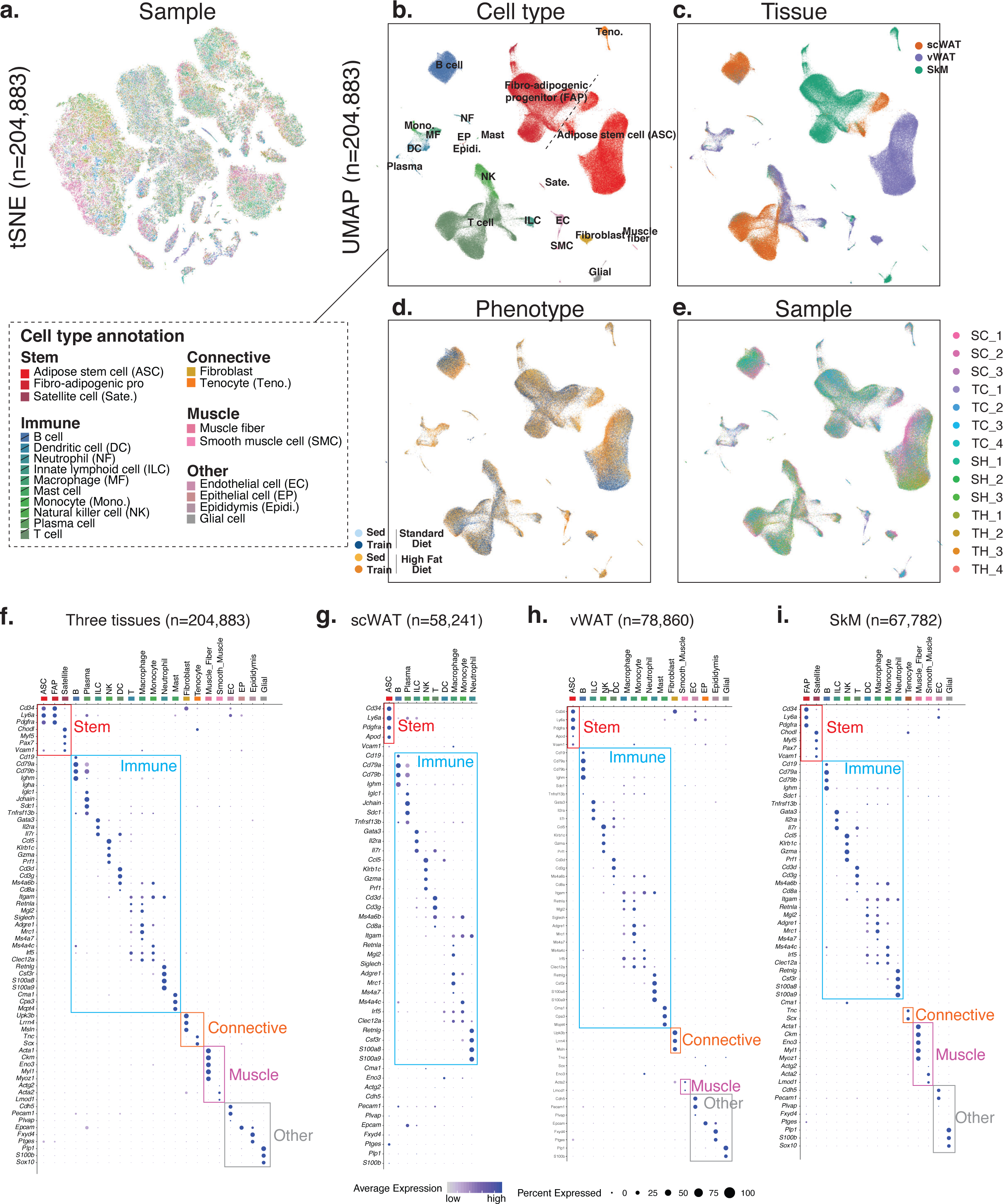
Different representations of single-cell atlas and cell-type-specific marker gene expression. **a-e,** Single-cell atlas of 204,883 cells across three tissues and four intervention groups. The tSNE (**a**) or UMAP (**c-e**) plot is coloured by sample (**a,e**), cell type (**b**), tissue (**c**), and intervention group (**d**). SC, sedentary chow (standard) diet; TC, training chow (standard) diet; SH, sedentary high-fat diet; TH, training high-fat diet. **f-i,** Cell-type-specific marker gene expression across the three tissues and in each individual tissue. A list of abbreviations used in this figure appear in the Methods.

**Extended Data Fig. 3:**
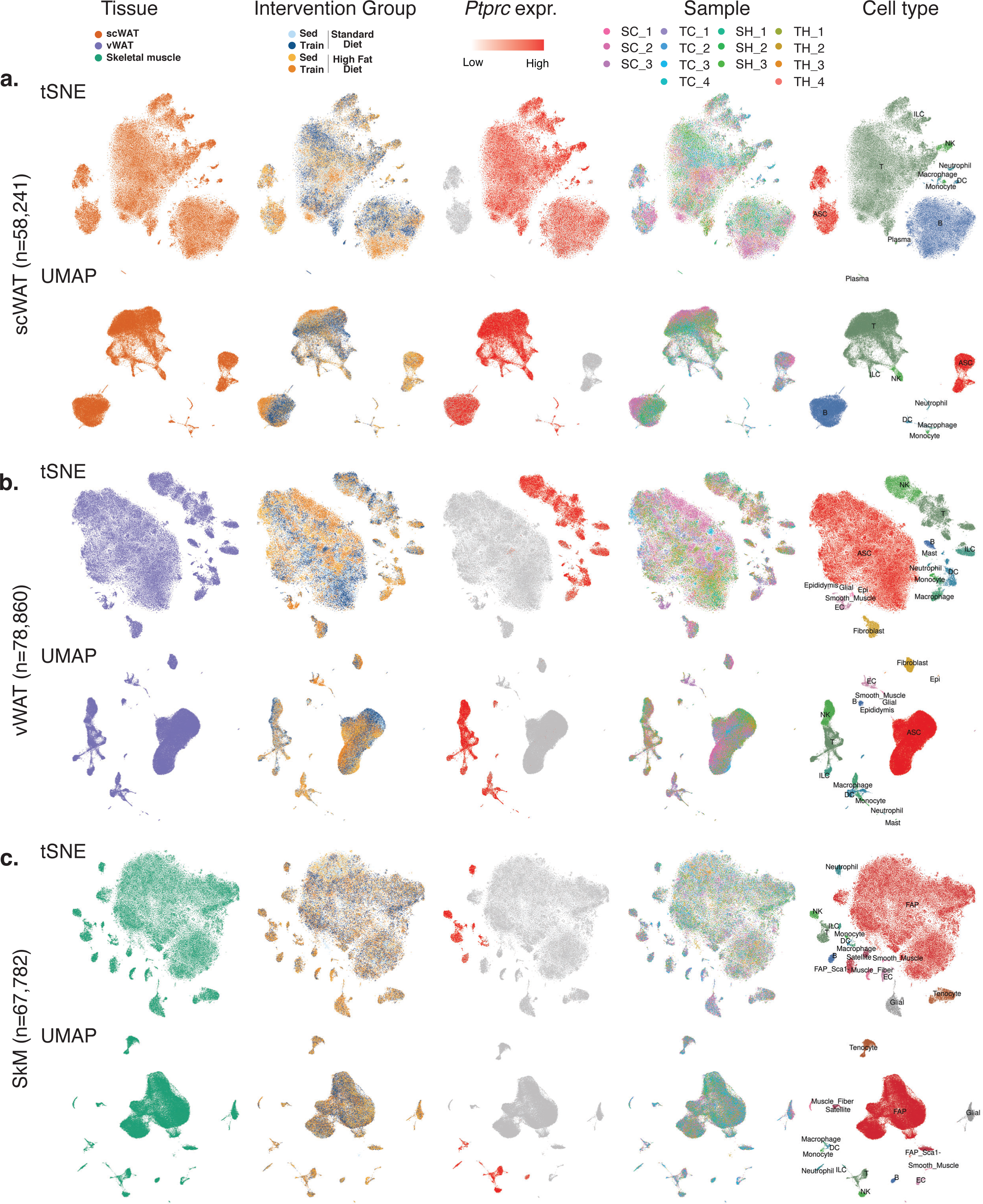
Different representations of single-cell maps for each tissue. **a-c**, Single-cell maps of scWAT (**a**), vWAT (**b**) and SkM (**c**) across four intervention groups. The tSNE and UMAP plots in each panel are coloured by tissue type, intervention group, *Ptprc* expression, sample, and cell type. A list of abbreviations used in this figure appear in the Methods.

**Extended Data Fig. 4:**
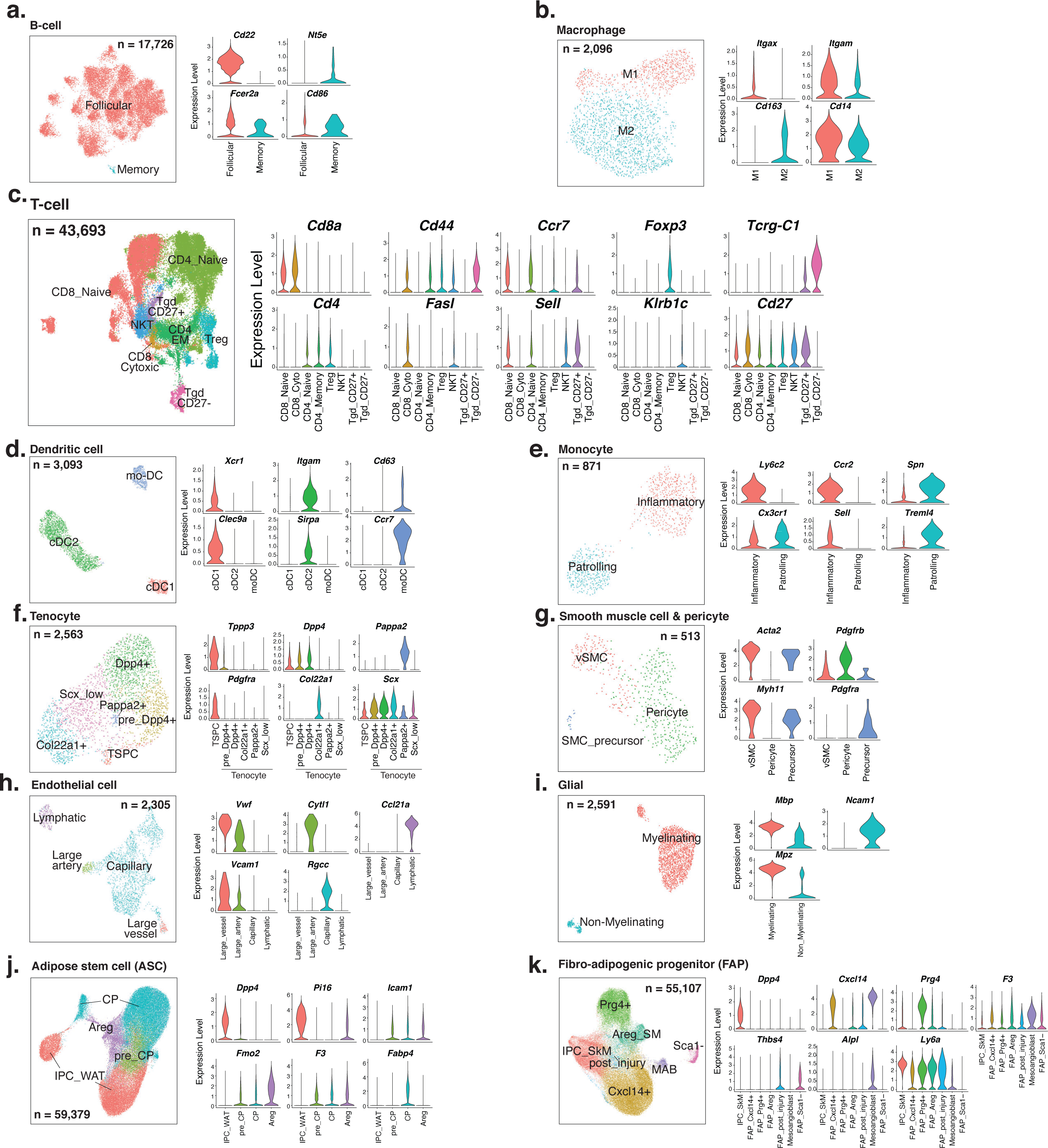
Sub-clustering of cell types across tissues. **a-k,** Cell subtypes or states of B-cell (**a**), macrophage (**b**), T-cell (**c**), dendritic cell (**d**), monocyte (**e**), tenocyte (**f**), smooth muscle cell (**g**), endothelial cell (**h**), glial cell (**i**), adipose stem cell (**j**), and fibro-adipogenic progenitor (**k**).

**Extended Data Fig. 5:**
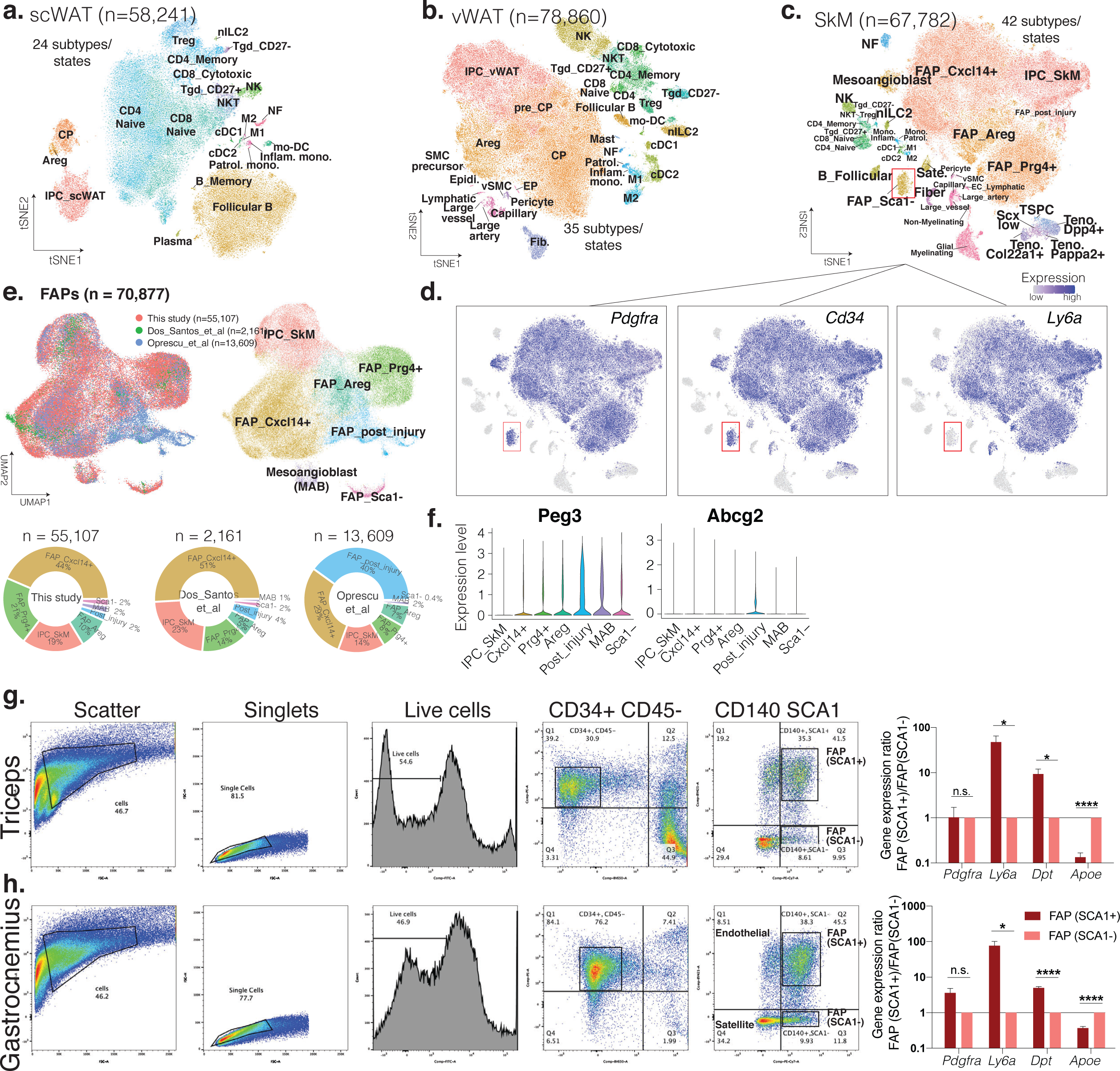
Single-cell subtype/state annotation in the three tissues and fibro-adipogenic progenitor characterization in skeletal muscle. **a-c,** Single-cell maps of scWAT (**a**), vWAT (**b**) and SkM (**c**) with cells coloured by cell subtype/state. **d,** Single-cell map of SkM with cells coloured by Pdgfra, Ly6a and Cd34 expression. **e,** Re-clustering of FAPs from this study with FAPs from two publicly available datasets. The cells are coloured by study (left) and cell state (right). Proportions of the seven distinct FAP states in each study are shown below. **f,** *Peg3* and *Abcg2* expression across FAP states in our single-cell data. **g-h,** FACS gating strategy and relative proportions of Sca1+ and Sca1-FAP populations in triceps (**g**) and gastrocnemius muscle (**h**). Statistical comparisons were performed using unparied t-test. P values are showns as *, < 0.05, ** < 0.01, *** < 0.001, **** < 0.0001. n.s., not significant. A list of abbreviations used in this figure appear in the Methods.

**Extended Data Fig. 6:**
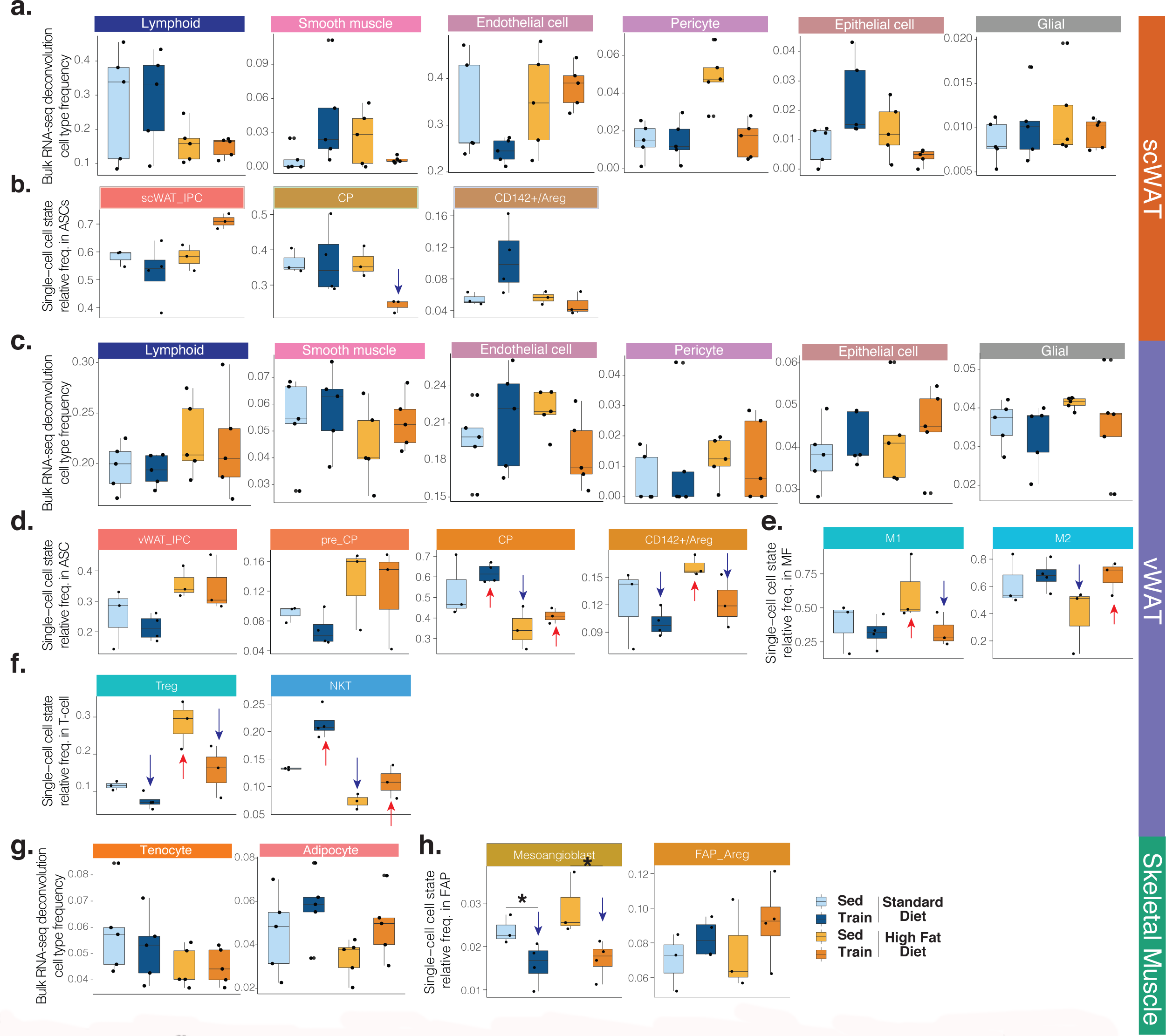
Cell type/state proportion changes with high-fat diet and exercise training. **a-b,** scWAT results. **a,** Sample-specific proportions of 6 cell types/groups after bulk RNA-seq data deconvolution across the four intervention groups. **b,** Sample-specific proportions of 3 states within ASCs based on our single-cell data in the four intervention groups. **c-f,** vWAT results. **c,** Sample-specific proportions of 6 cell types/groups after bulk ata deconvolution across the four intervention groups. **d-f,** Sample-specific proportions of 4 states within ASCs (**d**), 2 states of macrophages (**e**), and 2 states of T-cells (**f**) based on our single-cell data in the four intervention groups. **g-h,** Skeletal muscle results. **g,** Sample-specific proportions of 2 cell types after bulk RNA-seq data deconvolution across the four intervention groups. **h,** Sample-specific proportions of mesoangioblast and Areg_FAP within FAPs. A list of abbreviations used in this figure appear in the Methods.

**Extended Data Fig. 7:**
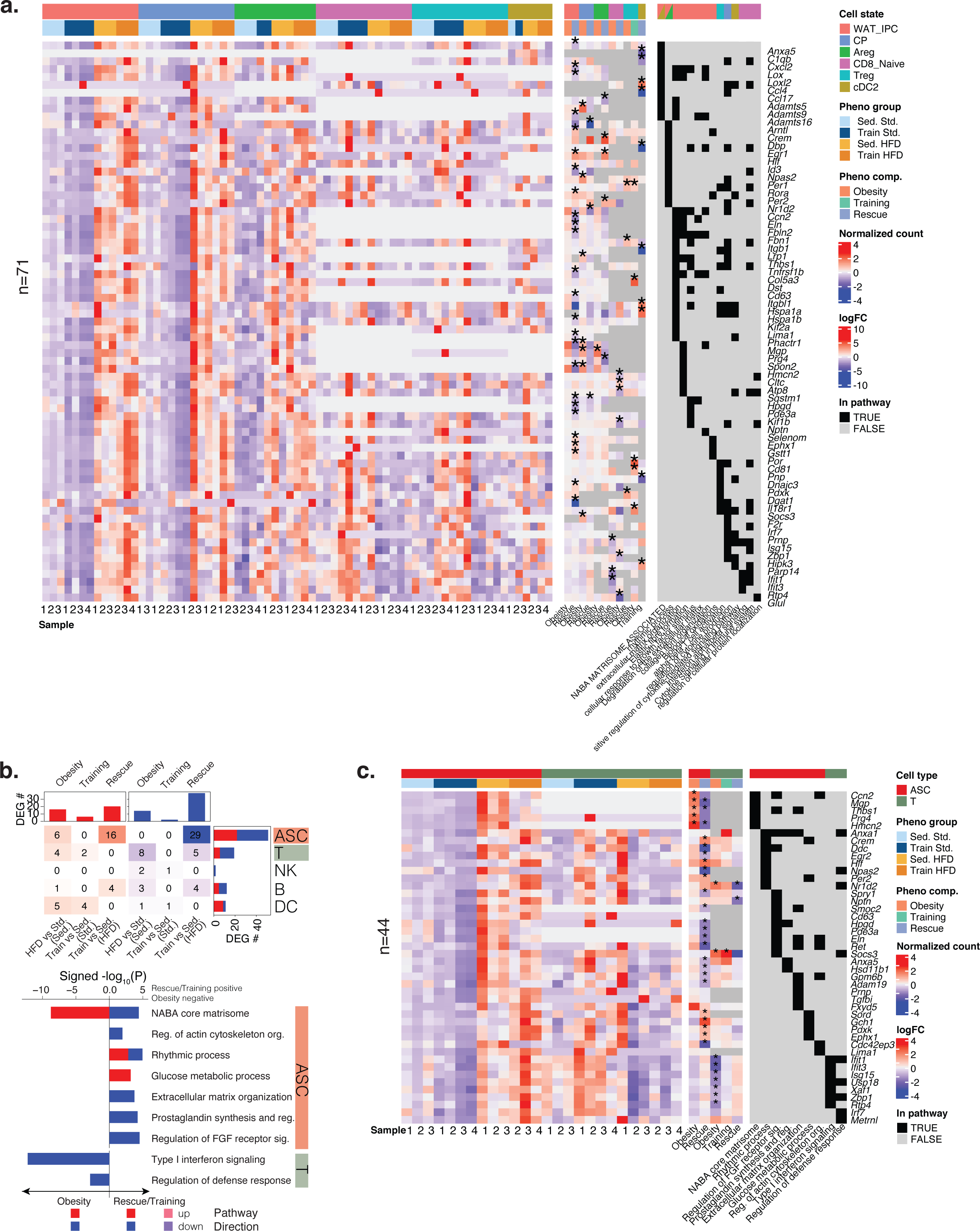
Cell-state and cell-type-specific transcriptomic changes with high-fat diet and exercise training in scWAT. **a,** Cell-state-specific DEGs and their membership in the significantly enriched pathways: sample-specific mean expression for each gene (left heatmap), their fold changes in the three comparisons whenever available (middle heatmap, * if significant), and their membership in the pathways (right heatmap). **b.** The number of cell-type-specific DEGs (heatmap) that are up-regulated (red) or down-regulated (blue) in our three comparisons. Pathways (bar plot) that are significantly enriched in cell-type-specific DEGs. **c,** Cell-type-specific DEGs and their membership in the significantly enriched pathways. The format is the same as in panel **a**. A list of abbreviations used in this figure appear in the Methods.

**Extended Data Fig. 8:**
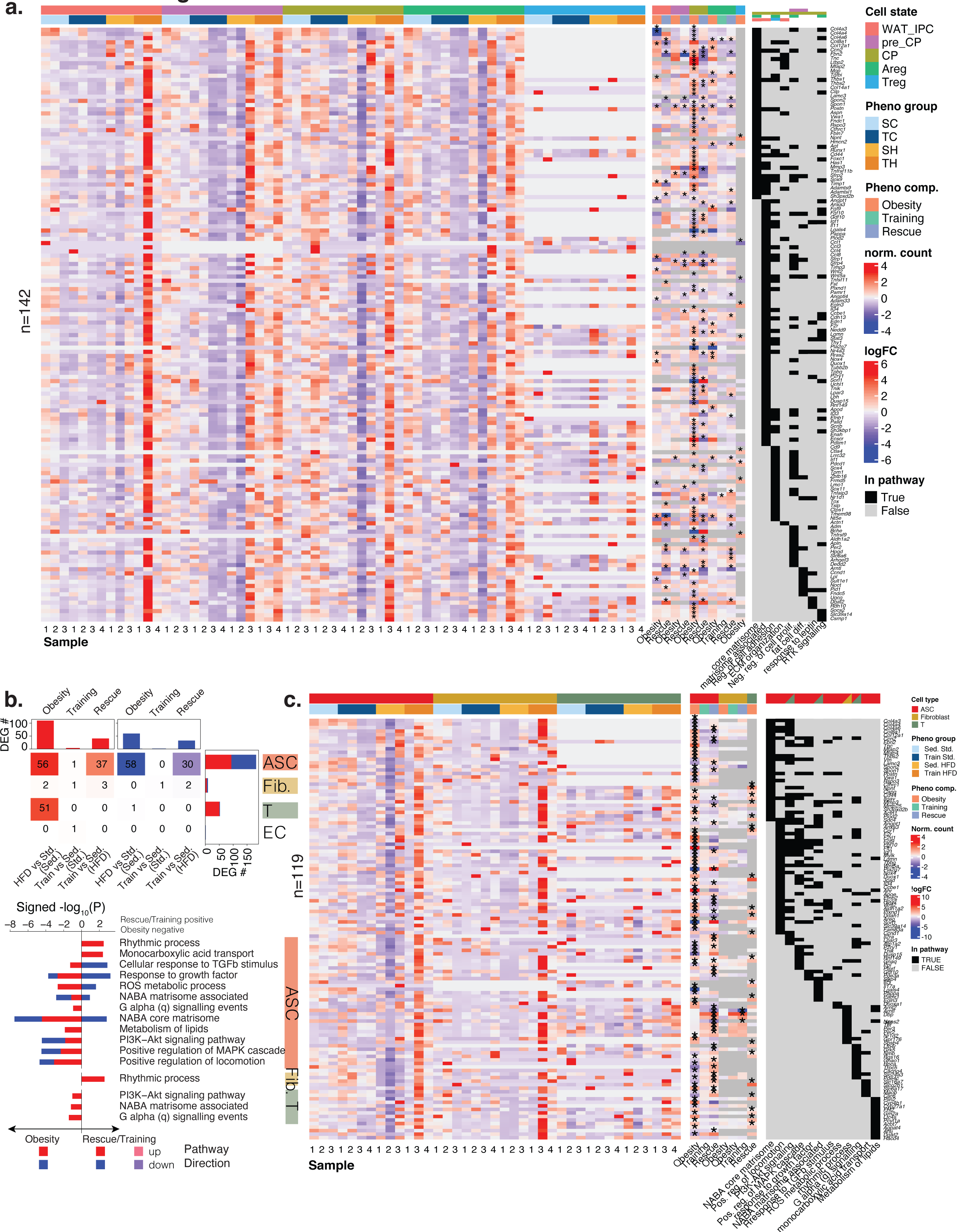
Cell-state and cell-type-specific transcriptomic changes with high-fat diet and exercise training in vWAT. **a,** Cell-state-specific DEGs and their membership in the significantly enriched pathways: sample-specific mean expression for each gene (left heatmap), their fold changes in the three comparisons whenever available (middle heatmap, * if significant), and their membership in the pathways (right heatmap). **b.** The number of cell-type-specific DEGs (heatmap) that are up-regulated (red) or down-regulated (blue) in our three comparisons. Pathways (bar plot) that are significantly enriched in cell-type-specific DEGs. **c,** Cell-type-specific DEGs and their membership in the significantly enriched pathways. The format is the same as in panel **a**. A list of abbreviations used in this figure appear in the Methods.

**Extended Data Fig. 9.**
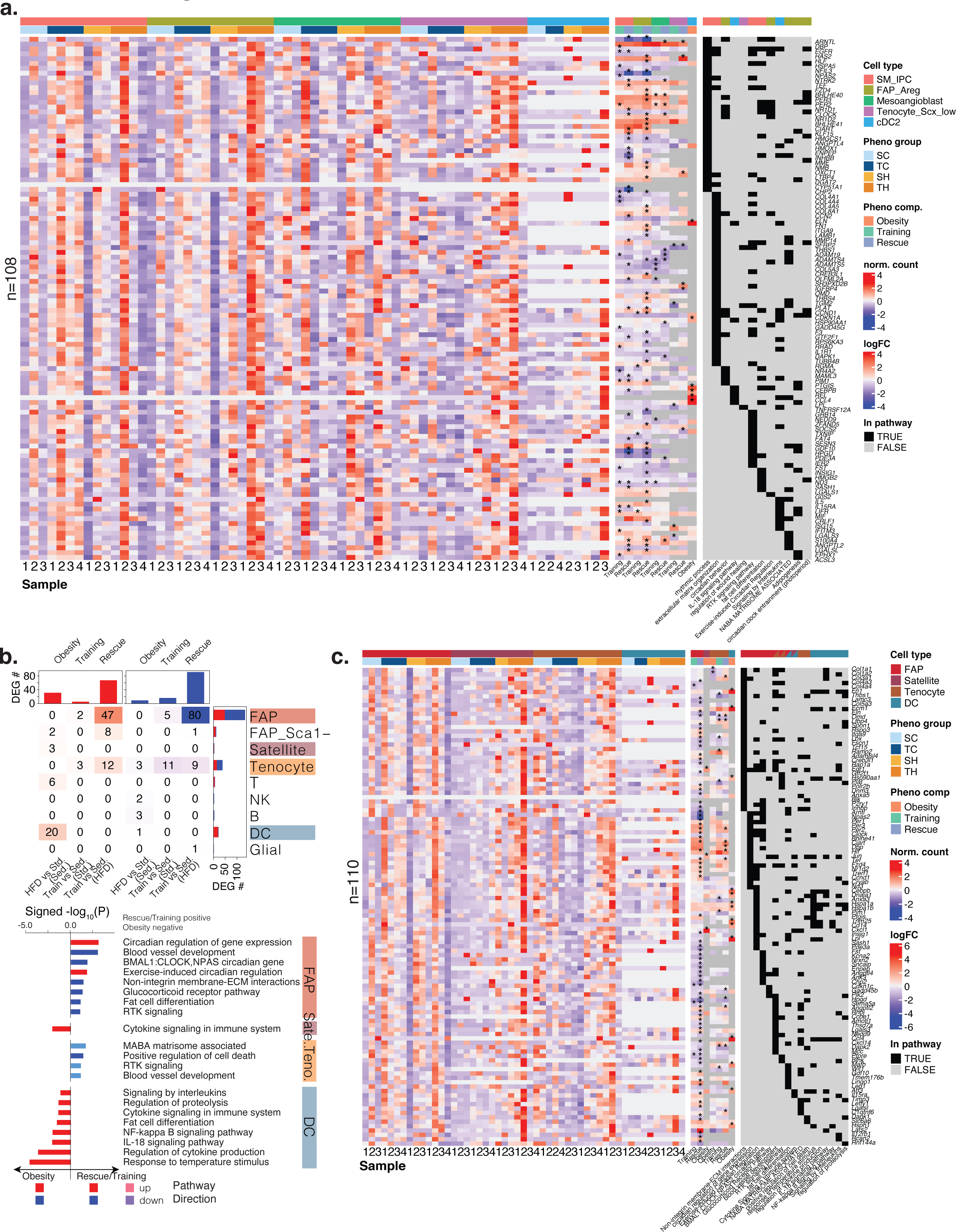
: Cell-state and cell-type-specific transcriptomic changes with high-fat diet and exercise training in SkM. **a,** Cell-state-specific DEGs and their membership in the significantly enriched pathways: sample-specific mean expression for each gene (left heatmap), their fold changes in the three comparisons whenever available (middle heatmap, * if significant), and their membership in the pathways (right heatmap). **b.** The number of cell-type-specific DEGs (heatmap) that are up-regulated (red) or down-regulated (blue) in our three comparisons. Pathways (bar plot) that are significantly enriched in cell-type-specific DEGs. **c,** Cell-type-specific DEGs and their membership in the significantly enriched pathways. The format is the same as in panel **a**. A list of abbreviations used in this figure appear in the Methods.

**Extended Data Fig. 10:**
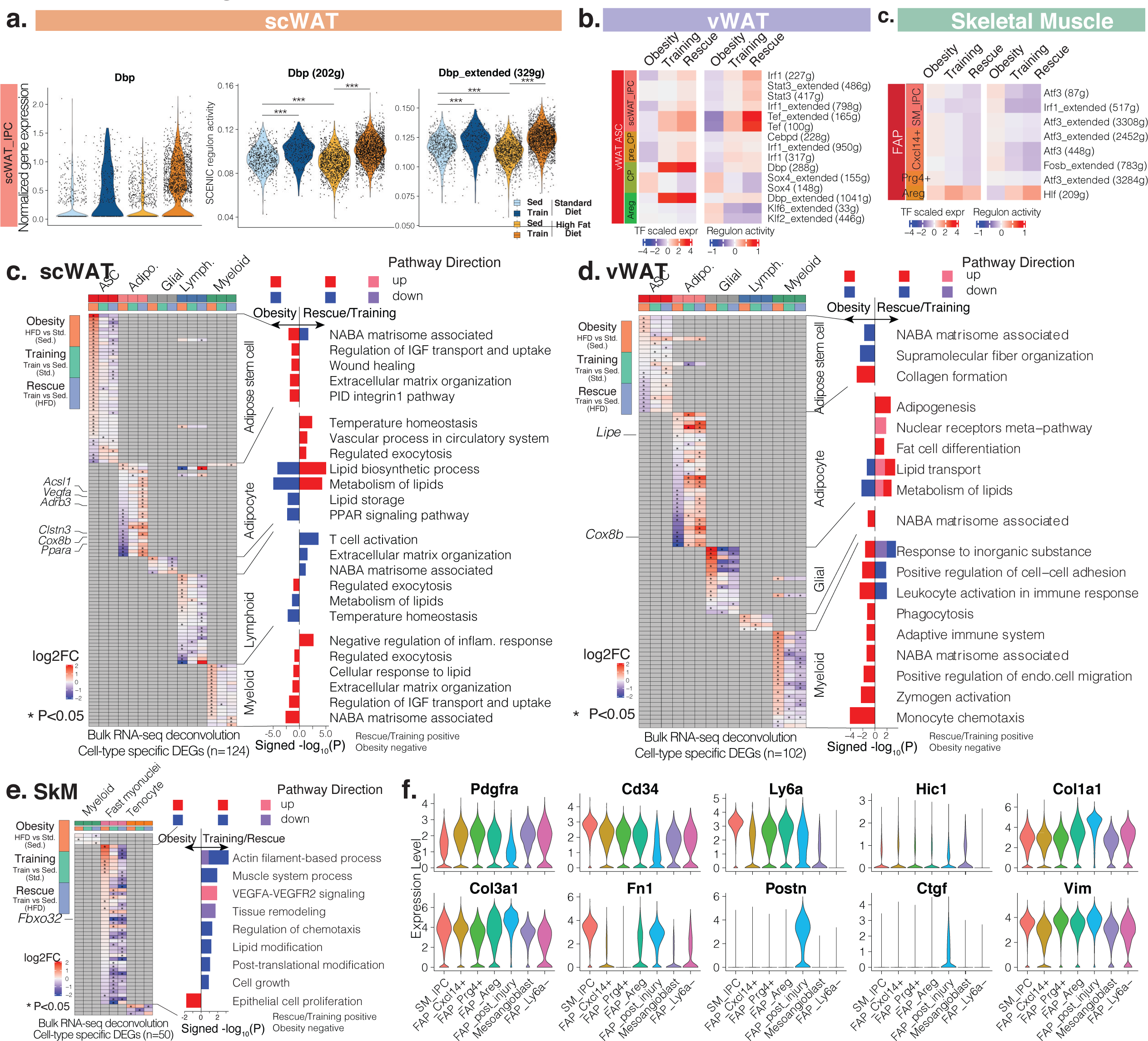
Regulon activity and deconvolved transcriptomic changes. **a.** Intervention group-specific Dbp expression and regulon activity in IPCs of scWAT. The number of genes regulated by Dbp is in parentheses. ***, p <0.001. **b-c.** Cell state-specific TF expression (left heatmap) and regulon activity (right heatmap) changes with the three comparisons in vWAT (**b**) and SkM (**c**). Only TF and regulon with p value less than 0.05 are shown. The number of genes regulated by each TF is in parentheses. **c-e,** After deconvolution cell group-specific DEGs (heatmap) and pathways (bar plot) that are up-regulated or down-regulated in our three comparisons in scWAT (**c**), vWAT (**d**) and SkM (**e**). X-axis of the bar plot shows -log_10_p value with rescue/training pathways being positive, and obesity being negative. The bars are coloured by pathway direction in the three comparisons (red/pink: up-regulated, blue/purple, down-regulated). **f.** Marker gene expression in Sca1+ and Sca1-FAPs. The markers are from a cell type similar to Sca1-FAP that is identified in heart. A list of abbreviations used in this figure appear in the Methods.

**Extended Data Fig. 11:**
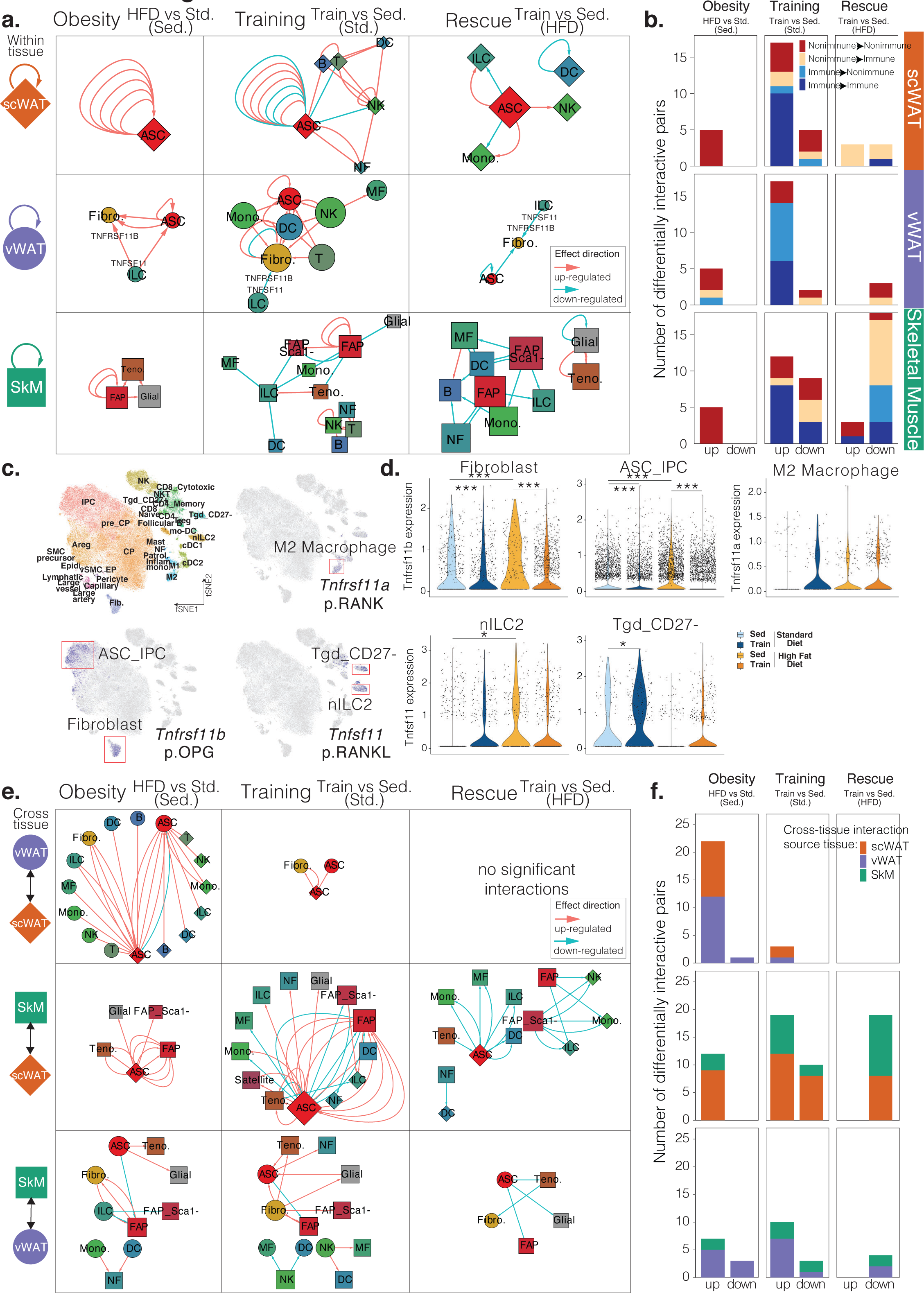
Within- and cross-tissue communication. **a,** Within-tissue ligand-receptor networks across the three tissues and three comparisons. Cell types (nodes) are shaped by tissue (diamond: scWAT, circle: vWAT, square: skeletal muscle) and sized by outdegree. Ligand-receptor interactions (edges) are directed from ligand to receptor, and coloured by effect direction (pink: up-regulated, blue: down-regulated). **b,** The number of differentially interactive ligand-receptor pairs that are up- and down-regulated across the three tissues and three comparisons at cell-type level. Each bar is coloured by if the ligand or the receptor is from immune or non-immune cell type. **c,** vWAT single-cell map with cells coloured by cell subtype/state, or expression of *Tnfsf11*, *Tnfrsf11a*, or *Tnfrsf11b*. **d,** Cell subtype/state-specific*Tnfsf11*, *Tnfrsf11a*, and *Tnfrsf11b* expression in the four intervention groups. * p < 0.05, *** p < 0.001. **e,** Cross-tissue ligand-receptor networks across three pairs of tissues and three comparisons. The nodes and edges are formatted the same as in panel **a**. **f,** The number of differentially interactive ligand-receptor pairs that are up- and down-regulated across three pairs of tissues and three comparisons. Each bar is coloured by tissue source of the ligand. A list of abbreviations used in this figure appear in the Methods.

**Extended Data Fig. 12:**
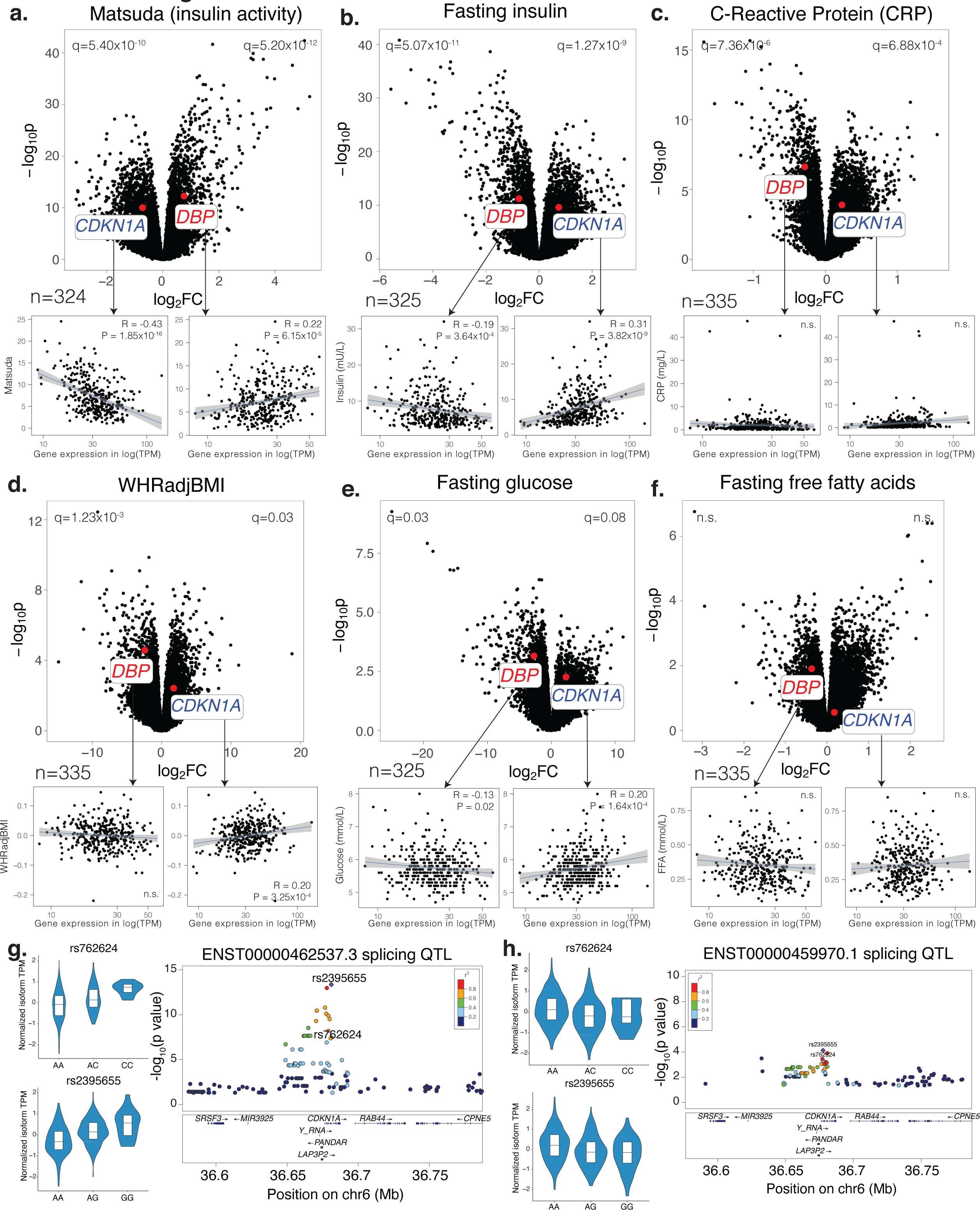
Association of DBP and CDKN1A with metabolic traits, and CDKN1A splicing QTLs in human. **a-f,** *DBP* and *CDKN1A* association with matsuda, an indicator of insulin activity (**a**); fasting insulin (**b**); CRP, an indicator of inflammation (**c**); waist hip ratio adjusted for BMI (**d**); fasting glucose (**e**); fasting free fatty acids (**f**) in scWAT of METSIM subjects. Genes (dots in upper plots) and subjects (dots in lower plots) are plotted **g-h,** Association of two SNPs (rs762624 and rs2395655) in *CDKN1A* with one transcript isoform (**g**) but not another (**h**) in METSIM. Violin plots show transcript levels associated with genotypes. Locus plots show splicing QTL p values and r-square values with nearby variants.TPM, reads mapped to transcript per million mapped reads.

**Supplementary Fig. 1:**
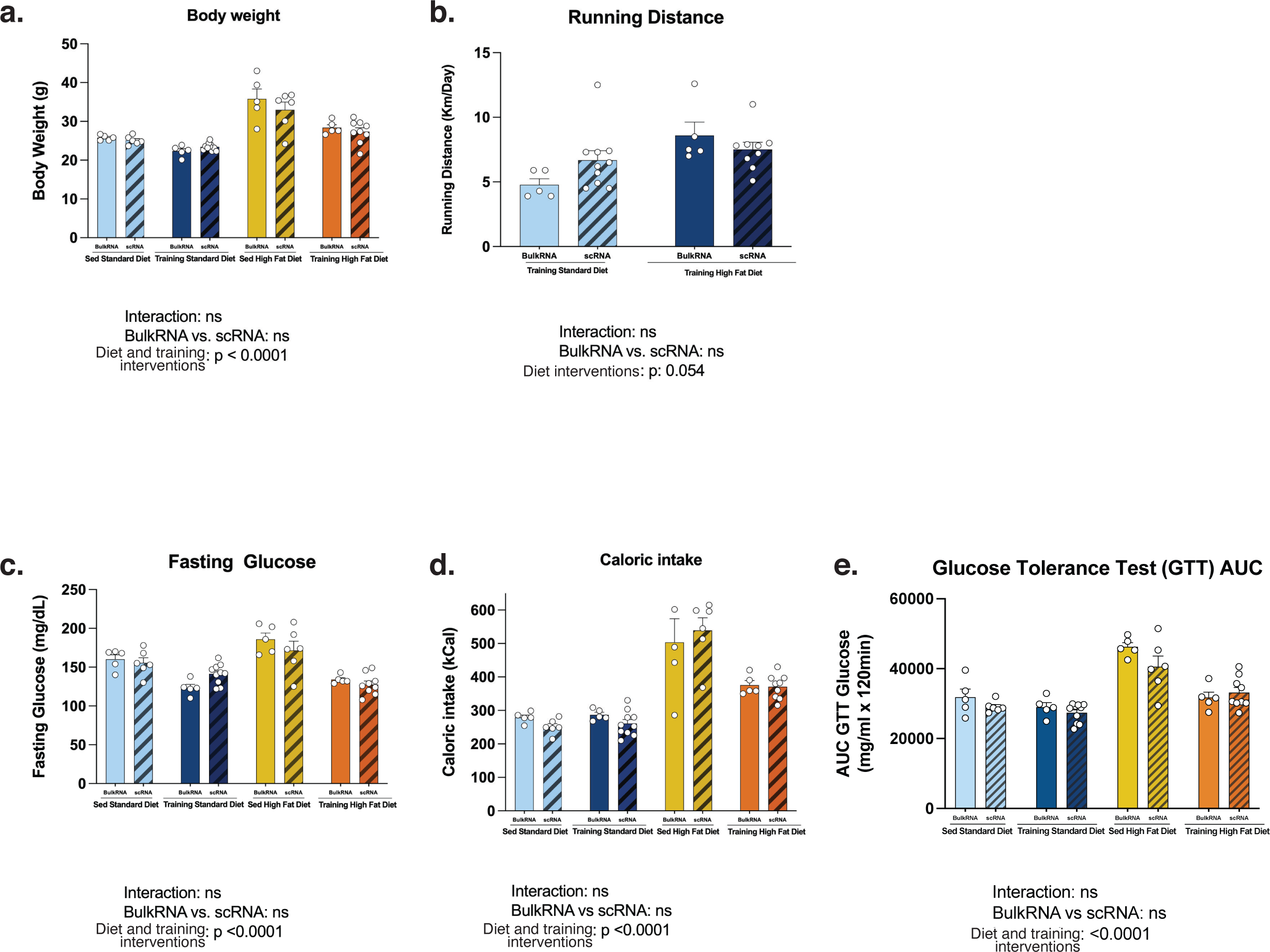
Phenotypic responses of the bulk and single-cell mouse cohorts. **a-e,** Body weight (**a**), running distance (**b**), fasting glucose (**c**), caloric intake (**e**), and glucose tolerance test (GTT) result for mice used in bulk and single-cell analysis across the four intervention groups. There is no statistically significant difference between the two cohorts for the meatrics shown. Statistical comparisons were performed using two way ANOVA. ns, not significant; AUC, area under curve. Other abbreviations used in this figure appear in the Methods.

**Supplementary Fig. 2:**
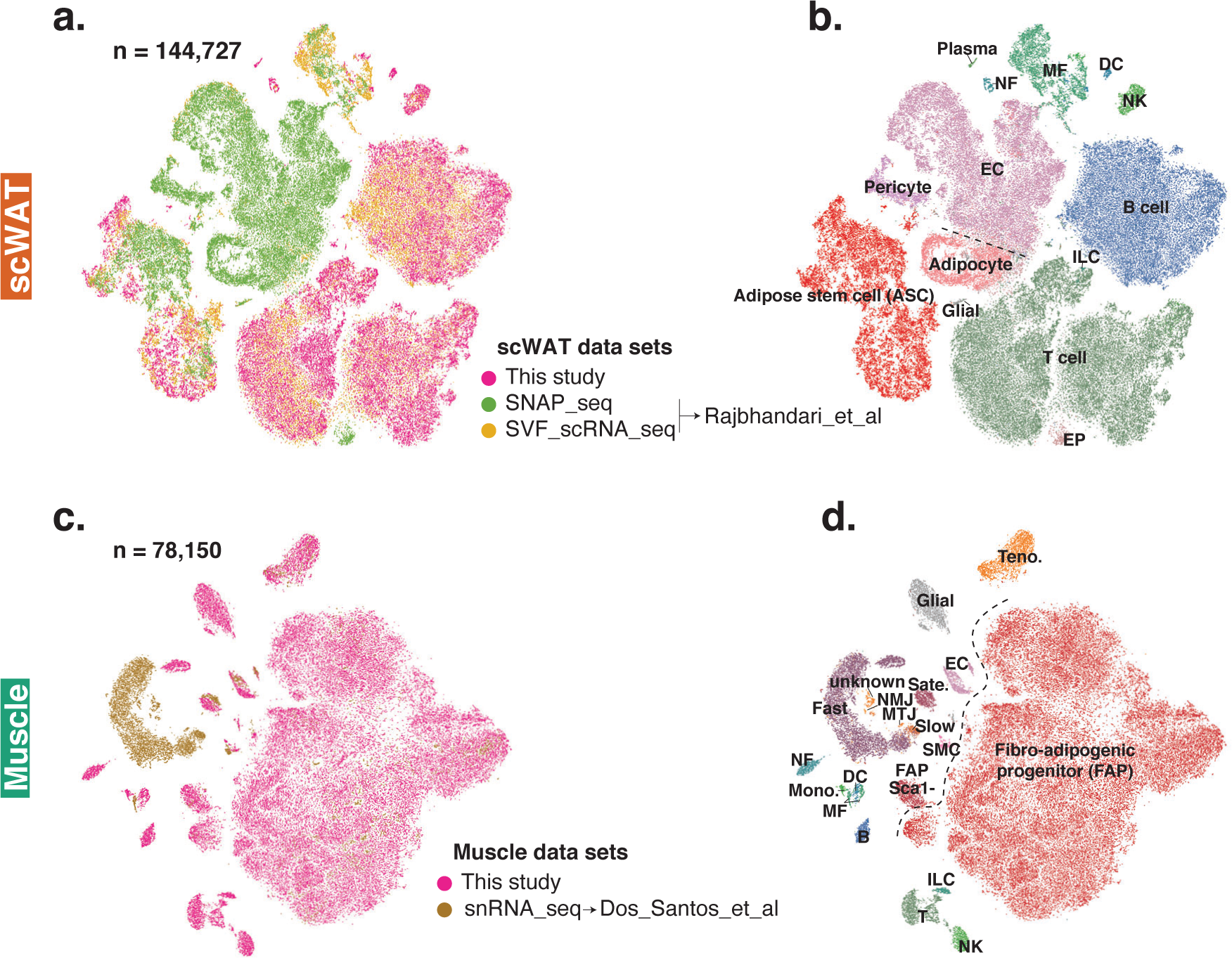
Integration of this study with tissue-matched publicly available single-cell studies. **a-b,** Single-cell clustering of 144,727 cells from this study and GSE133486. The UMAP plot is coloured by study (**a**) and cell type (**b**). **c-d,** Single-cell clustering of 78,150 cells from this study and GSE150065. The UMAP plot is coloured by study (**c**) and cell type (**d**). A list of abbreviations used in this figure appear in the Methods.

**Supplementary Fig. 3:**
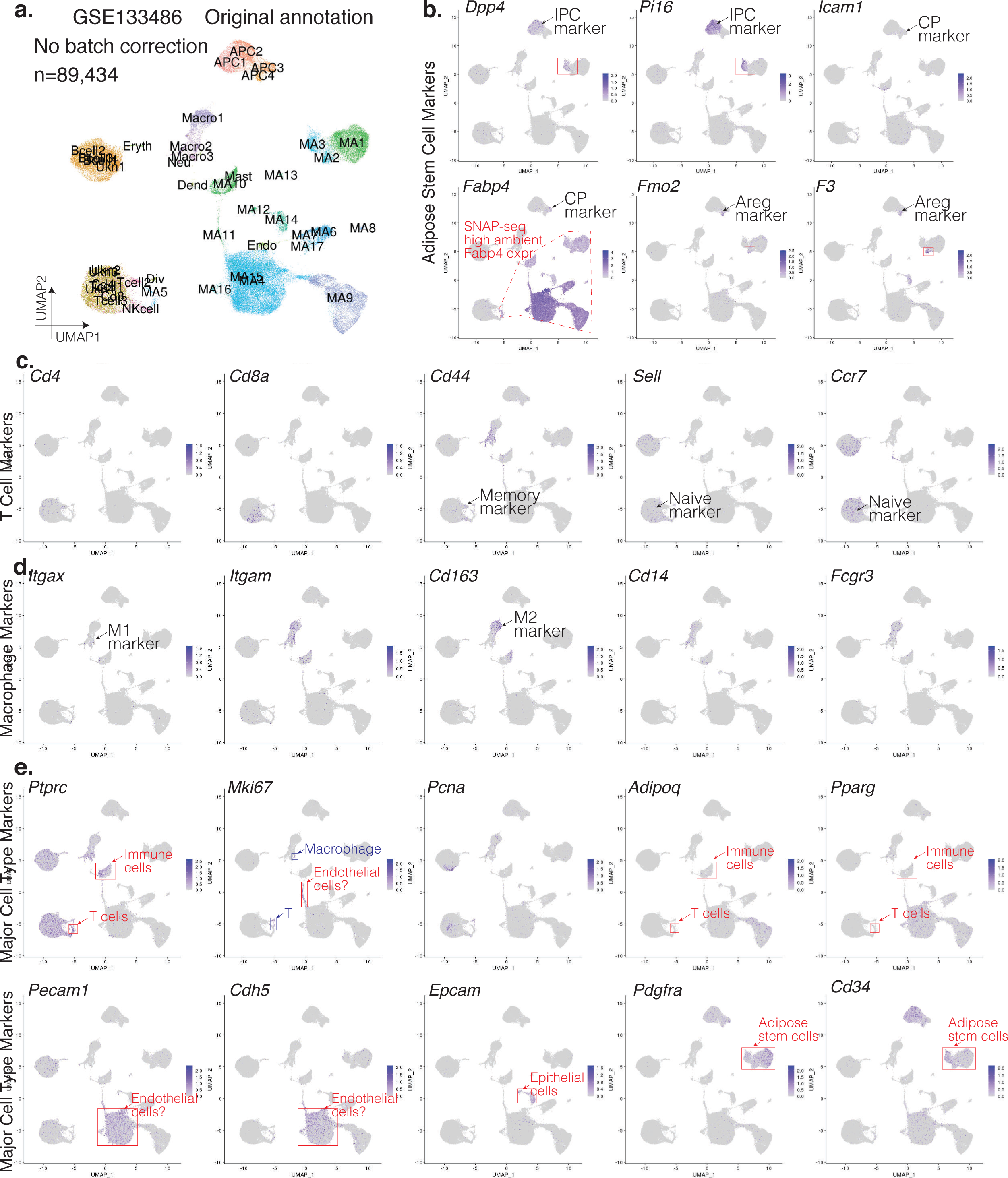
Original cell type annotation and marker gene expression for a single-cell mouse subcutaneous white adipose tissue study. **a,** Single-cell clustering of 89,434 cells in GSE133486. The UMAP plot is coloured by original cell type annotation provided by the authors. MA, mature adipocyte; APC, adipose progenitor cell; Dend, dendritic cell; Macro, macrophage; Neu, neutrophil; Endo, endothelial cell; Div, dividing cell; Eryth, erythrocyte; Ukn, unknown. **b-f,** Marker gene expression for adipose stem cell (**b**), T-cell (**c**), macrophage (**d**), and other major cell types (**e**). Cell clusters with uncertain marker gene expression are boxed. A list of abbreviations used in this figure appear in the Methods.

**Supplementary Fig. 4:**
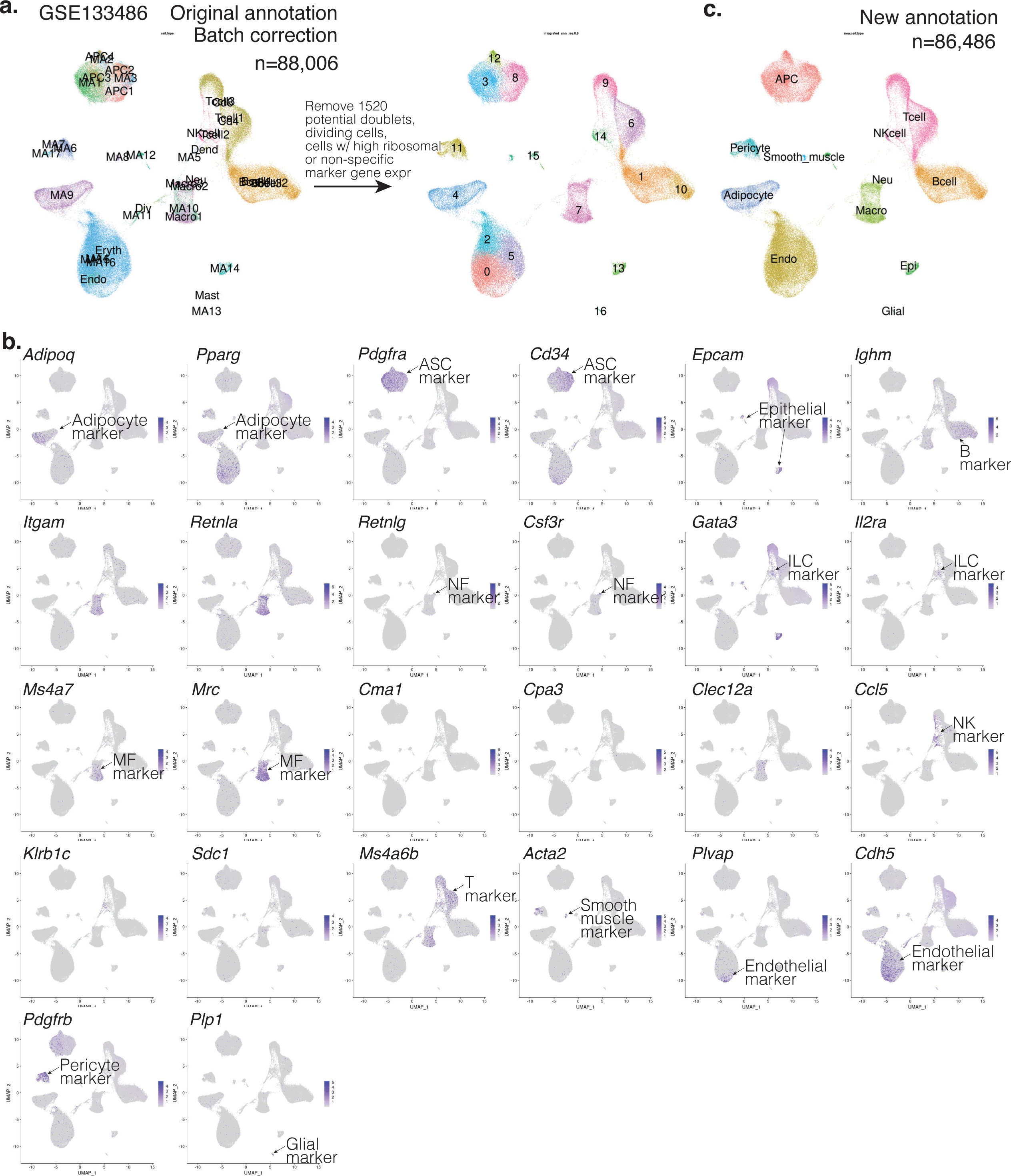
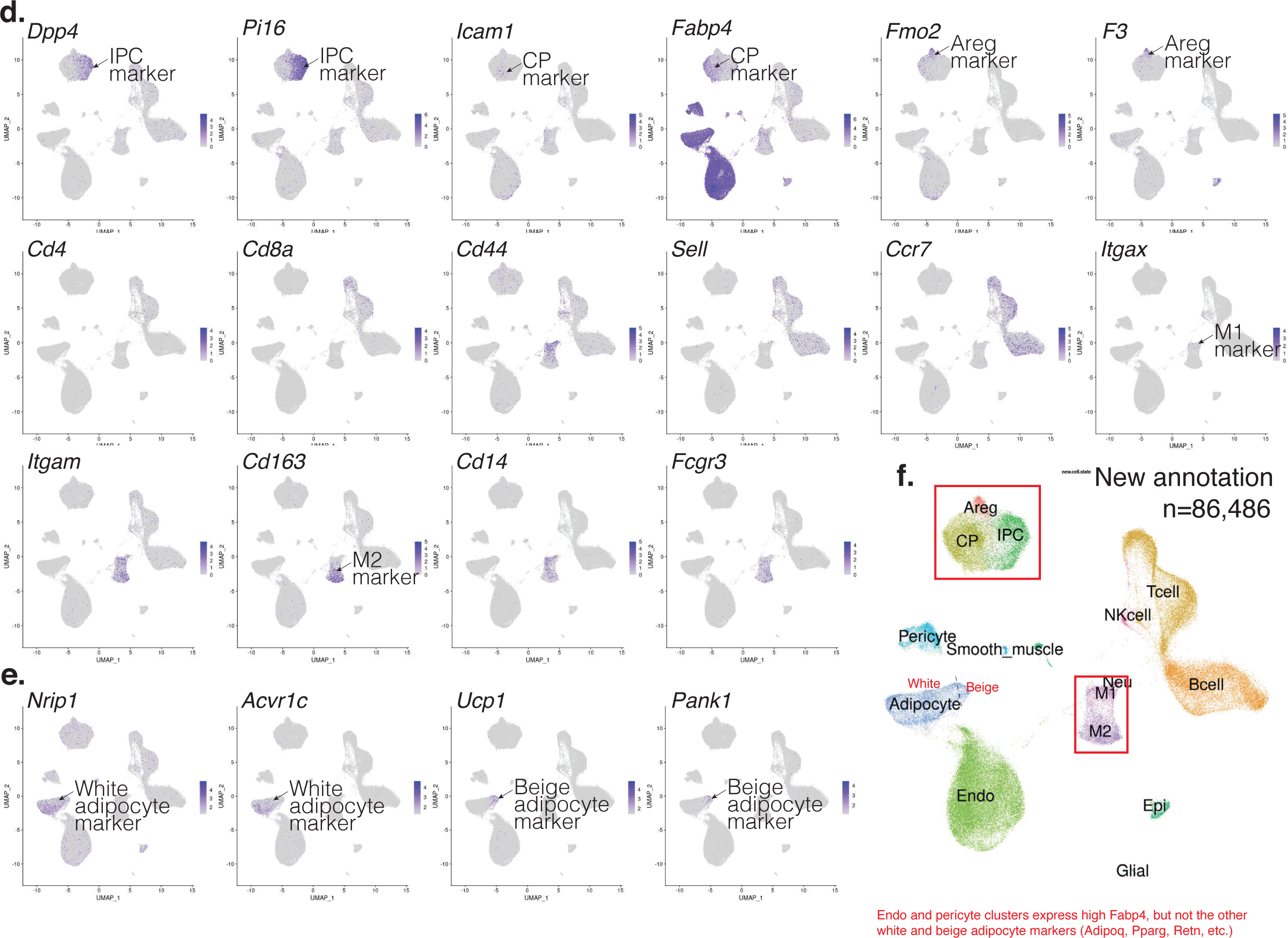
New cell type and state annotation and marker gene expression for GSE133486. **a,** QC, batch correction, and de novo clustering of the single-cell data set. **b,** Cell-type marker gene expression. **c,** New cell type labels assigned to each cluster after QC and batch correction. **d,** Cell-state marker gene expression. **e,** White and beige adipocyte marker gene expression. **f,** New cell state labels. A list of abbreviations used in this figure appear in the Methods.

**Supplementary Fig. 5:**
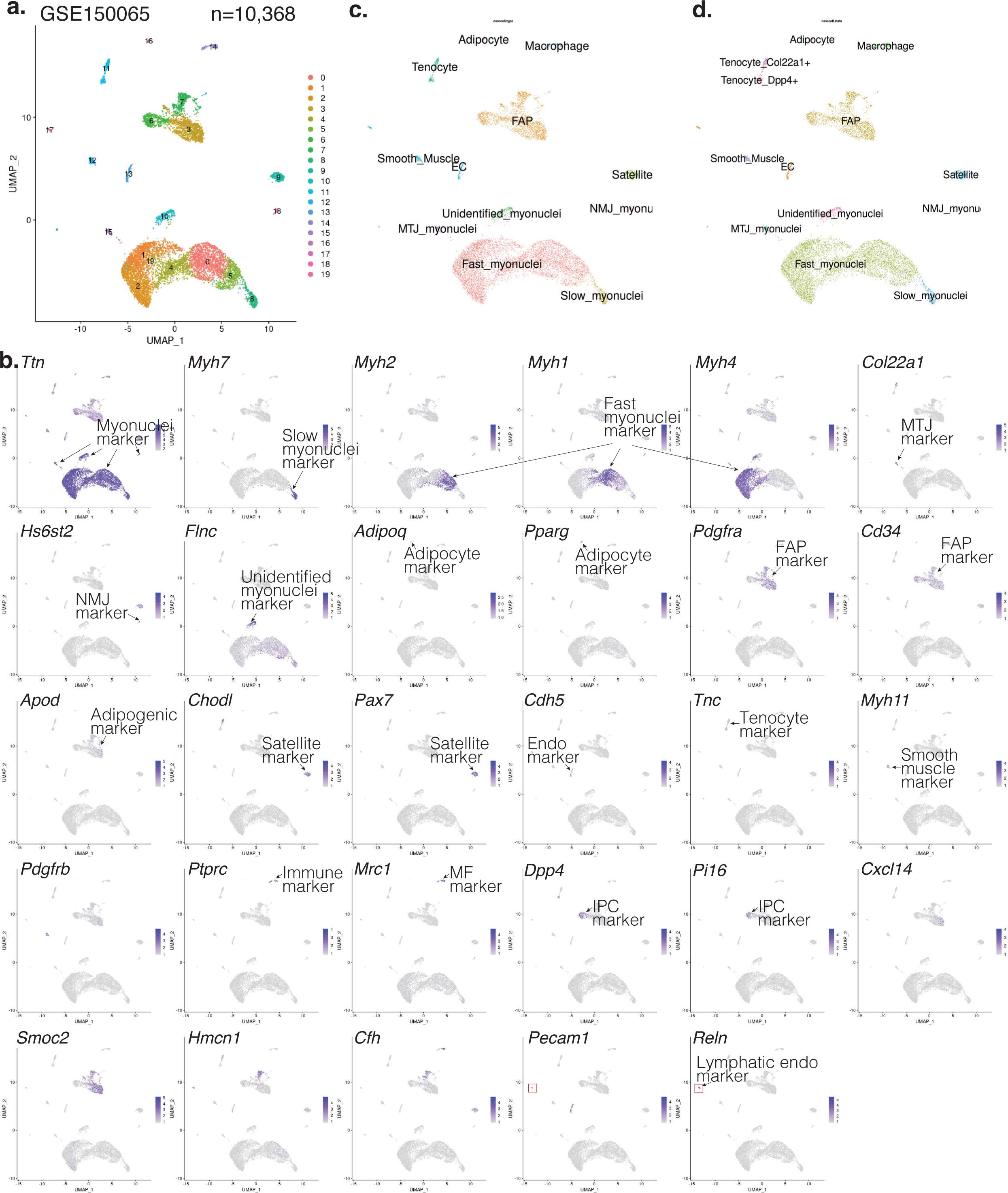
Original cell type annotation and marker gene expression for a single-cell mouse skeletal muscle study. **a,** Single-cell clustering of 10,368 cells in GSE150065. The UMAP plot is coloured by clusters. **b,** Cell-type marker gene expression. **c,** Same UMAP plot coloured by original cell type labels provided by the authors. **d,** Further split of tenocytes into two states. MTJ, myotendinous junction specialization; NMJ, neuromuscular junction. Other abbreviations used in this figure appear in the Methods.

**Supplementary Fig. 6:**
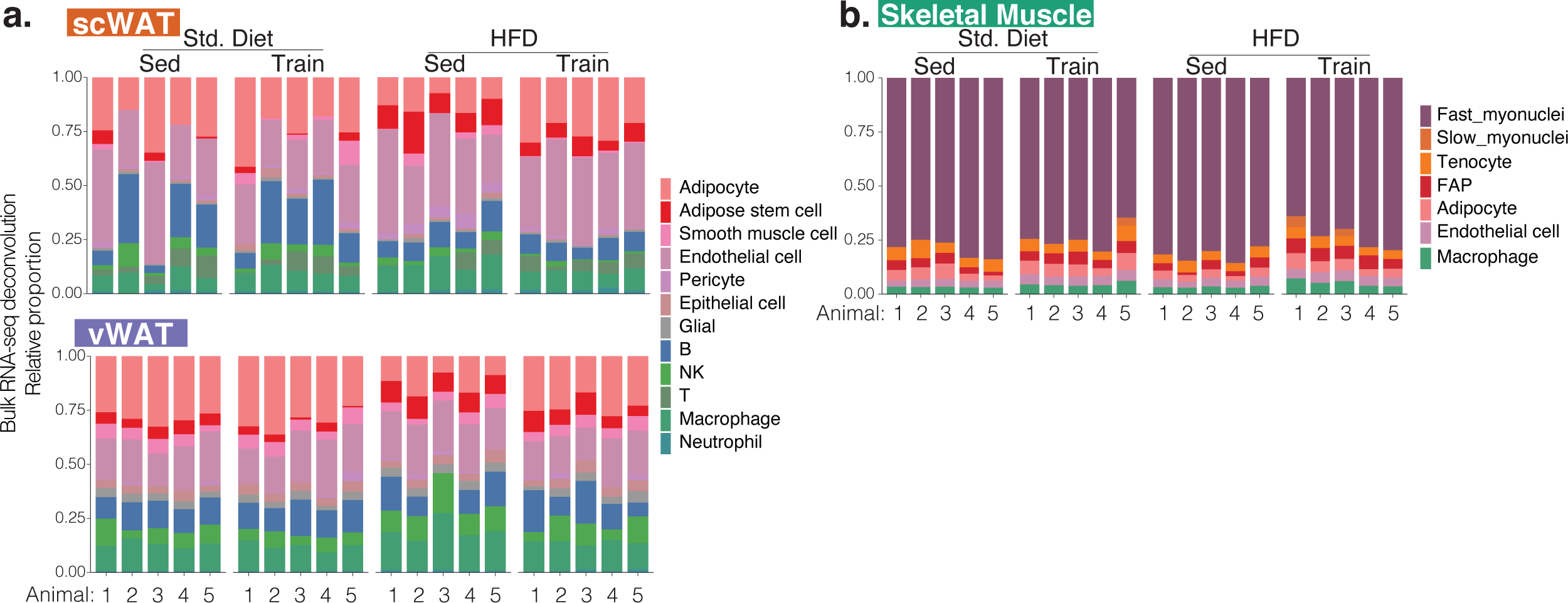
Sample-specific deconvolution. **a-b,** Relative cell-type proportions in each sample after deconvolving bulk RNA-seq data from two WAT depots (**a**) and skeletal muscle (**b**). Each bar is coloured by cell type.

**Supplementary Fig. 7:**
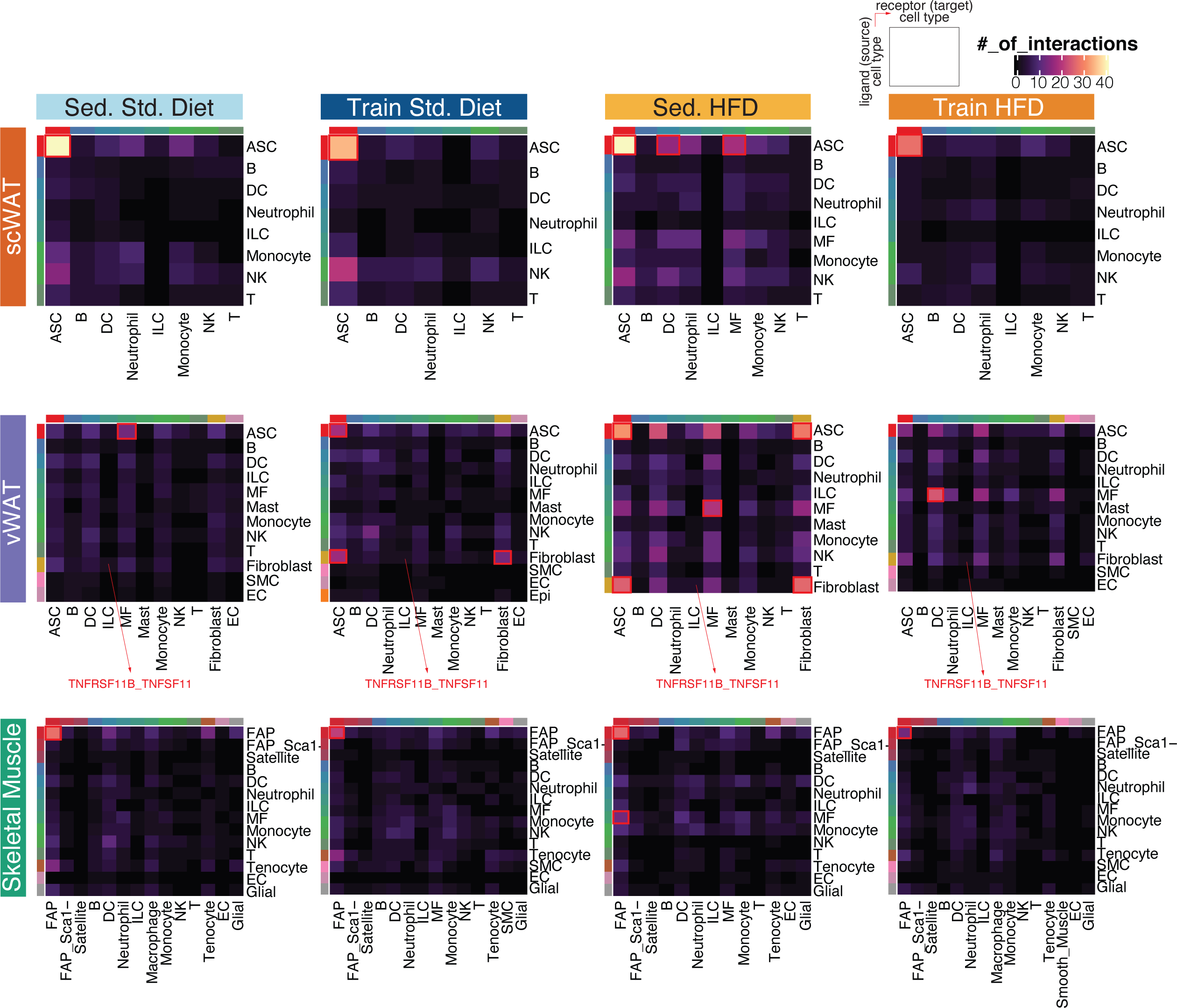
Within-tissue communication at cell-type level. Within-tissue ligand-receptor networks across the three tissues (row panels) in the four intervention groups (column panels). Number of interactions (heatmaps) between source cell types with ligand (rows) and target cell types with interacting receptors (columns) are plotted. Cell-type pairs with relatively higher number of interactions in each heatmap are highlighted using red rectangles. A list of abbreviations used in this figure appear in the Methods.

**Supplementary Fig. 8:**
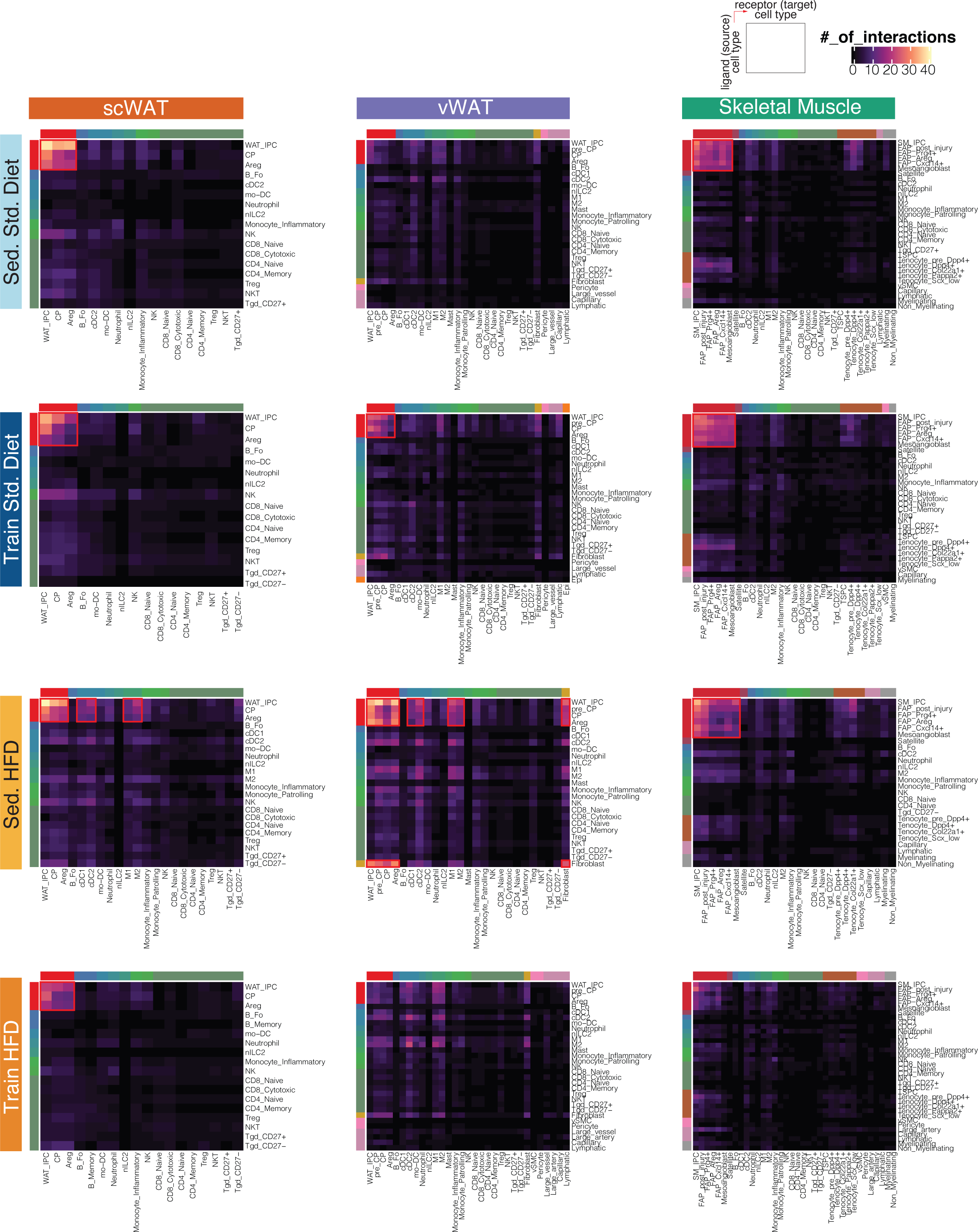
Within-tissue communication at cell-state level. Within-tissue ligand-receptor networks across the three tissues (column panels) in the four intervention groups (row panels). Number of interactions (heatmaps) between source cell states with ligand (rows) and target cell states with interacting receptors (columns) are plotted. Cell-state pairs with relatively higher number of interactions in each heatmap are highlighted using red rectangles. A list of abbreviations used in this figure appear in the Methods.

**Supplementary Fig. 9:**
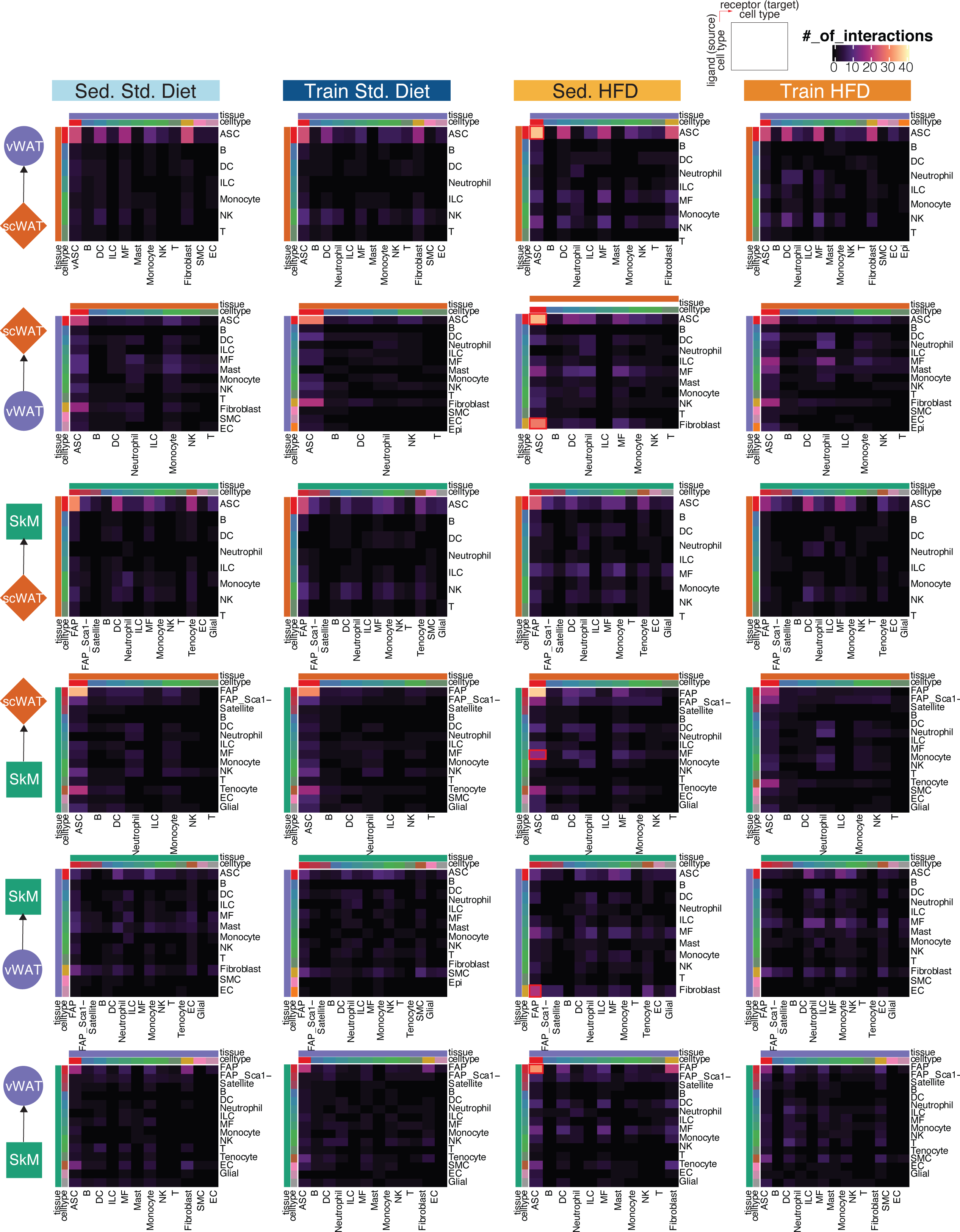
Cross-tissue communication at cell-type level. Cross-tissue ligand-receptor networks across tissue pairs (row panels) in the four intervention groups (column panels). Number of interactions (heatmaps) between source-tissue cell types with ligand (rows) and target-tissue cell types with interacting receptors (columns) are plotted. Cell-type pairs with relatively higher number of interactions in each heatmap are highlighted using red rectangles. A list of abbreviations used in this figure appear in the Methods.

**Supplementary Fig. 10:**
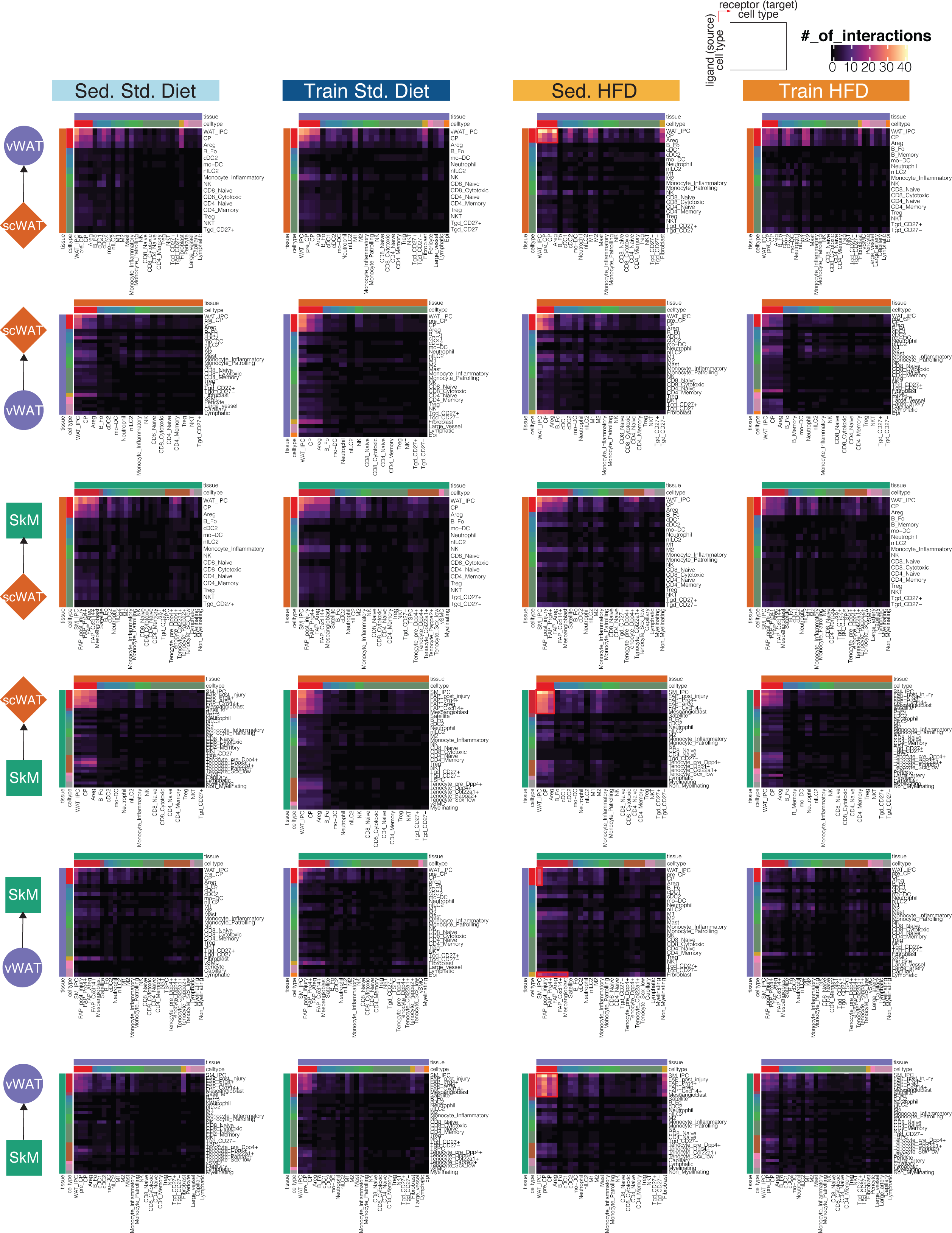
Cross-tissue communication at cell-state level. Cross-tissue ligand-receptor networks across tissue pairs (row panels) in the four intervention groups (column panels). Number of interactions (heatmaps) between source-tissue cell states with ligand (rows) and target-tissue cell states with interacting receptors (columns) are plotted. Cell-state pairs with relatively higher number of interactions in each heatmap are highlighted using red rectangles. A list of abbreviations used in this figure appear in the Methods.

## Notes

### Competing Interest Statement

The authors have declared no competing interest.

